# Dietary fiber intake associates with improved survival and microbiome composition in allogeneic hematopoietic cell transplantation

**DOI:** 10.64898/2026.06.03.729896

**Authors:** Jenny Paredes, Tyler Funnell, Peter Adintori, Anqi Dai, Natalie Smith, Ruben J. Faustino Ramos, Preet Kaur, Zeyang Li, Khyati Pathak, Jennifer Funes, Kristen Victor, Romina Ghale, Nathan Doung, Jennifer M. Haber, Keimya Sadeghi, Charlotte Pohl, Ashley Huang, Luigi A. Amoretti, Ariel Molina, Mirae Baichoo, Harold Elias, Oriana Miltiadous, Anastasia I. Kousa, Andri L. Lemarquis, Scott James, Ana Catarina Gradissimo de Oliveira, Urvi A. Shah, Patrick Pirrotte, Justin Cross, Jonathan U. Peled, Marina Burgos da Silva, Teng Fei, Marcel RM van den Brink

## Abstract

Diet is linked to changes in gut microbiota and metabolite production with clinical relevance in several disease settings, although these effects remain poorly defined. We performed prospective, real-time diet monitoring (37,929 food items, 3,837 patient days) and longitudinal microbiome and metabolite profiling (1,230 fecal samples) in a clinical cohort of 173 patients undergoing allogeneic hematopoietic cell transplantation. Patients with pre-transplant fiber intake above the cohort average had significantly improved overall survival (p=0.014) and reduced incidence of grades 2–4 acute graft-versus-host disease (GVHD) (p=0.032) post-transplant. Those consuming insoluble fiber had increased microbial diversity, enriched butyrate-producing taxa, and depleted *Enterococcus*. Those who developed lower gastrointestinal GVHD had reduced fecal butyrate levels. In a GVHD preclinical model, we confirmed that a fiber-enriched diet increased survival, cecal butyrate, and regulatory-to-conventional T cell ratio. Thus, we demonstrated that dietary fiber has clinical significance as a modifiable factor with microbiome-mediated effects.

## Introduction

Despite known causal links between diet and disease, evidence-based dietary guidance for patients undergoing cancer therapies remains limited. Recent studies have found that dietary patterns strongly shape gut microbiome composition and function, and that changes in the gut microbiome have downstream effects on cancer treatment response and toxicity^1,2,3,4,5,6^. High-fiber diets are of particular interest given promising evidence across multiple diseases,^5,7,8,9,10^ including after cancer immunotherapy^6,11^, and demonstrated cardiometabolic benefits^12^. Fiber is also associated with beneficial gut microbial diversity. A recent study reported that a high-fiber diet intervention outperformed fecal microbiota transplantation in repairing antibiotic-induced gut dysbiosis^13^, which is characterized by reduced diversity, depletion of commensal anaerobes such as *Blautia*, and expansion of facultative aerobes such as *Enterococcus*^14,15,16^.

Mechanistic studies investigating the gut microbiome have demonstrated that microbial metabolites from fiber fermentation, particularly short-chain fatty acids (SCFAs) such as butyrate, acetate, and propionate, have immunoregulatory and protective effects on the intestinal epithelium, including reinforcing intestinal barrier integrity and suppressing pro-inflammatory signaling^17,18^. For instance, butyrate enhances regulatory T-cell differentiation and limits Th1/Th17-mediated inflammation, mechanisms relevant to immune-mediated pathogenesis^19,20,21^. Bile acids are also microbial metabolites playing important roles in modulating inflammation and maintaining gut barrier integrity^22,23^.

Allogeneic hematopoietic cell transplantation (allo-HCT), a curative therapy for hematologic malignancies, provides a robust setting for evaluating the clinical relevance of dietary fiber and the microbiome. Conditioning regimens, antibiotics, and immunosuppressive therapies during allo-HCT induce gut dysbiosis, which correlates with graft-versus-host disease (GVHD) severity and transplant-related mortality^16,24,25,26,27^. This dysbiosis is characterized by a loss of diversity, decreased abundance of *Blautia,* and increased abundance of *Enterococcus*^15,16^. Despite evidence linking microbiome composition with these outcomes^27,28^, the association between dietary fiber intake and allo-HCT clinical endpoints has not been systematically evaluated. We therefore analyzed a prospective cohort of 173 allo-HCT recipients by collecting detailed dietary data daily in the clinical setting, corrected for portion sizes and nutritional intake in grams, and performing microbiome profiling. Patients with pre-transplant fiber intake above the cohort average had improved overall survival (OS), reduced incidence of severe acute GVHD, and an improved microbiota profile. In a GVHD preclinical model used to corroborate our clinical findings, we found that a fiber-enriched diet led to increased survival, increased cecal butyrate, and a higher regulatory-to-conventional T cell ratio.

## Results

### Fiber intake is positively associated with improved survival and lower risk of severe acute GVHD

Given prior work suggesting that dietary fiber intake can modulate the microbiome composition^1,2,3,4,5^, we hypothesized that high pre-transplant intake of fiber would be associated with improved OS and a lower incidence of severe acute GVHD in allo-HCT recipients. To test this hypothesis, we prospectively collected nutritional intake and clinical outcomes data from 173 recipients (**Table 1**). These nutritional intake data, recorded from day -7 to day +30 relative to transplant, encompass 37,929 individual food items (**Fig. 1a**) and 3,837 patient-days (**Fig. 1a**). We used *Computrition*, a hospital nutrition management software integrated with electronic health records, to document all served foods and portion sizes. Intake amounts were verified by dietitians and clinical research staff for each participant, and nutrient values were standardized using the USDA Food and Nutrient Database for Dietary Studies (FNDDS)^29^. In parallel, we collected serial fecal samples (average=7.3 samples per patient) from patients between day -7 and day 30 relative to transplant, totaling 1,230 samples (**Supplemental Fig. 1**).

**Figure 1.**
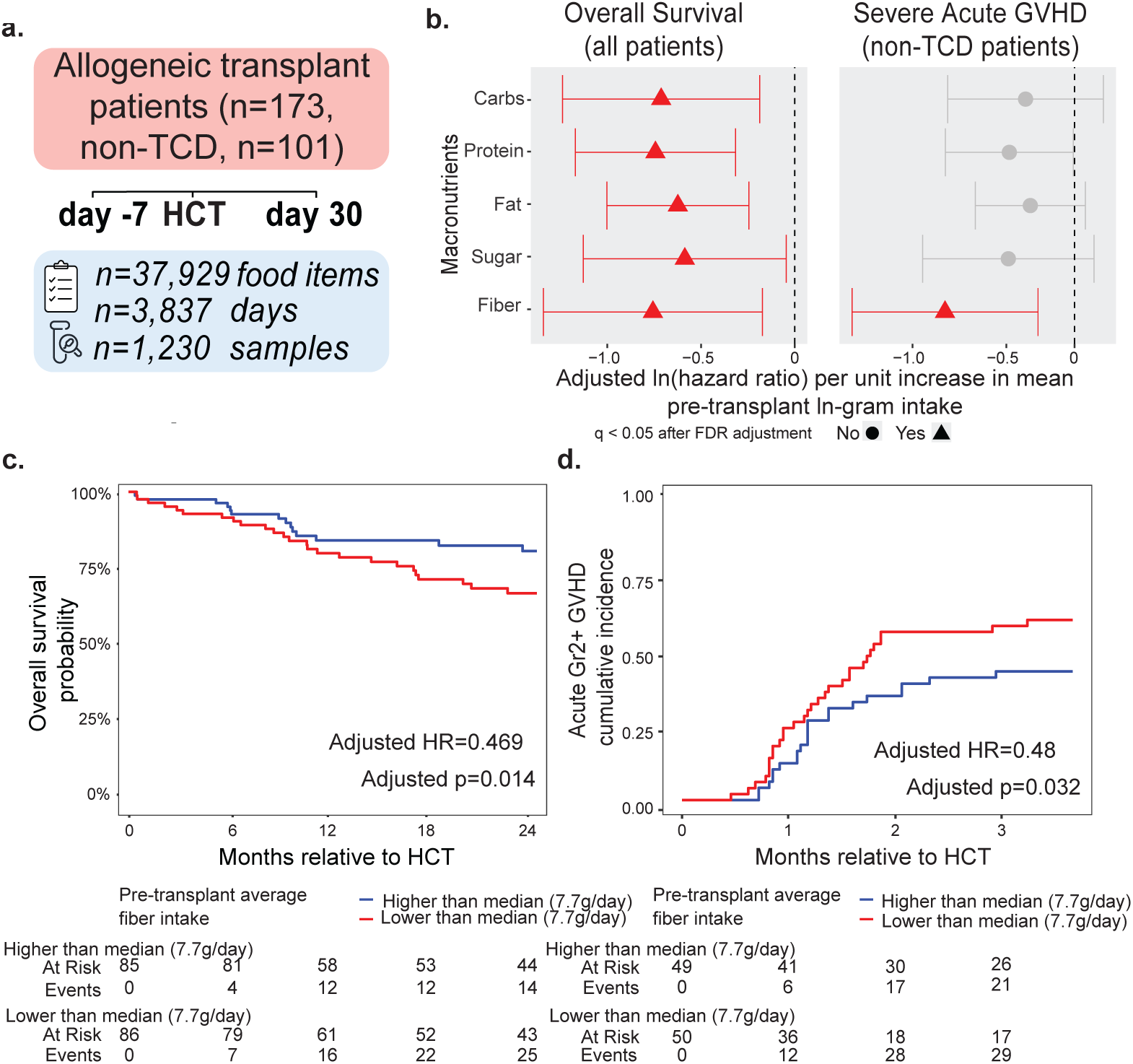
Higher average pre-transplant intake (ln-grams) of dietary fiber is associated with improved overall survival and reduced acute-GVHD following allo-HCT. a: Study design and nutritional data curation in 173 patients undergoing allogeneic hematopoietic cell transplantation (allo-HCT) included daily collection using the software *Compnutrition* followed by correction for portions eaten for every food item and calculation of macronutrients in grams. Non-TCD, non-T-cell depleted. Days: patient-days, data was recorded from dietary intake. Samples: fecal samples. b: Associations between overall survival (OS, of all patients, left [n=171]) and severe acute GVHD (aGVHD, of non-TCD patients, right [n=99]) with average ln-transformed pre-transplant (day -7 to day -1) intake (ln-grams/day) of macronutrients, carbohydrates, protein, fat, sugar, and fiber. Ln-transformed hazard ratios (HR) or sub-distributional HRs per unit increase in ln-gram intake were estimated for OS and aGVHD via multivariable Cox and Fine-Gray regression models, respectively. All models were adjusted for pre-transplant antibiotic exposure, disease type, conditioning intensity, and graft source (for OS models only). P-values were adjusted across macronutrients for each time-to-event outcome. Macronutrients for which FDR-adjusted p-values < 0.05 are shown with estimated log-HRs annotated by red triangles and red unadjusted 95% confidence intervals. No: not significant. Yes: significant. c: Kaplan-Meier estimates of OS by dietary fiber intake groups thresholded by the median ln-transformed pre-transplant intake (grams/day, n=171 patients with available pre-HCT dietary data). d: Aalen-Johansen estimates of severe aGVHD (grade ≥2) cumulative incidence in non-TCD patients by dietary fiber intake thresholded by the median pre-transplant intake (grams/day, n=99 patients with available pre-HCT dietary data).

**Supplemental Figure 1.**
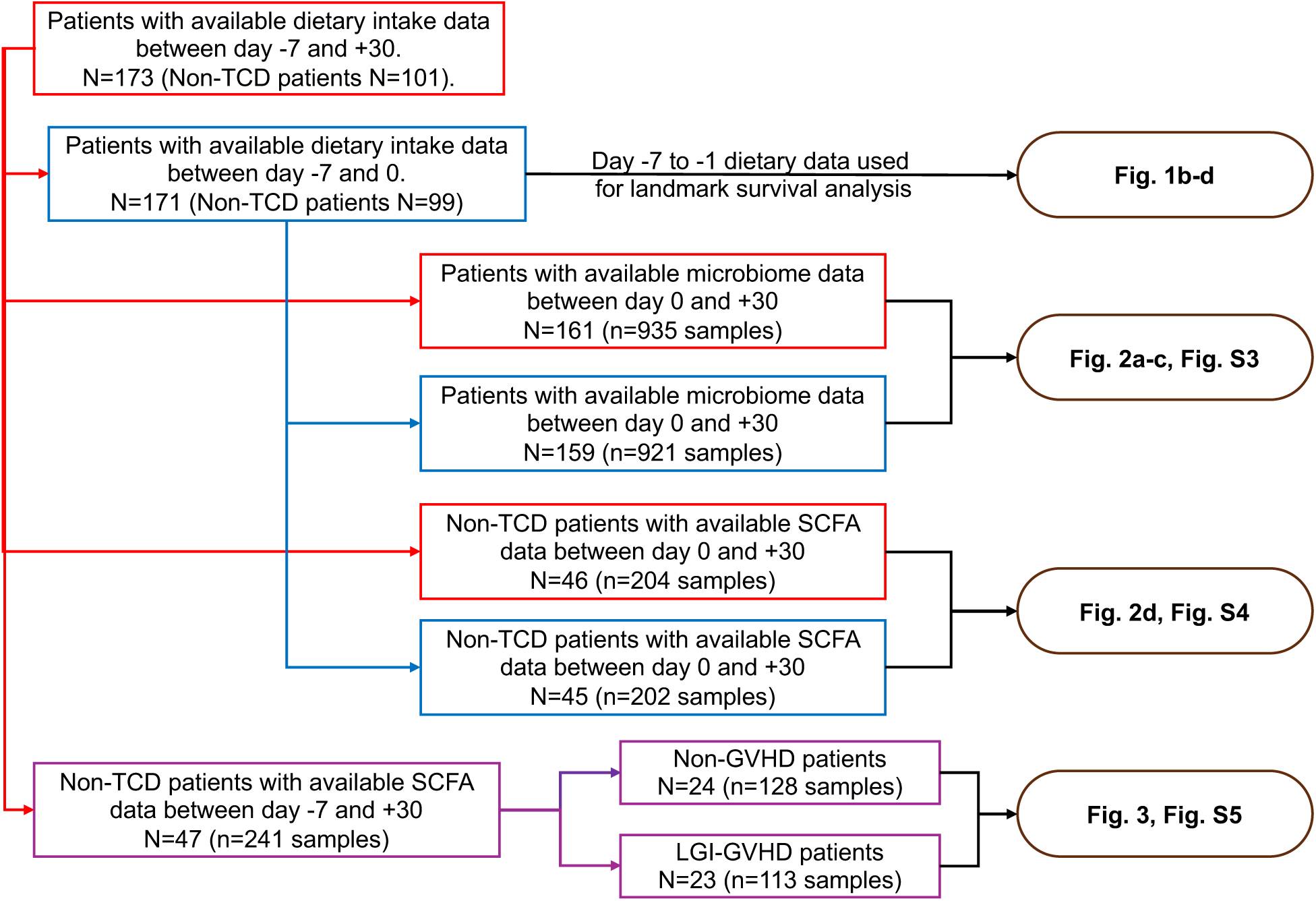
Data availability for dietary intake, microbiome, and SCFA analyses. Flowchart illustrating patient cohorts defined according to the availability of dietary intake data, microbiome sequencing (16S rRNA), and metabolite measurements across specified time windows. These subsets were used for survival analyses, microbiome profiling, and metabolite comparisons in subsequent figures.

**Table 1.** Characteristics of patients with available pre-HCT dietary data.

| Characteristic | N = 171<sup>1</sup> |
| --- | --- |
| **Age** | 60 (50, 66) |
| **Sex** |  |
| Male | 93 (54%) |
| Female | 78 (46%) |
| **Disease** |  |
| Acute myeloid leukemia | 67 (39%) |
| MDS/MPN | 57 (33%) |
| Non-Hodgkin's lymphoma | 17 (9.9%) |
| Acute lymphoid leukemia | 12 (7.0%) |
| Chronic lymphocytic leukemia | 2 (1.2%) |
| Myeloma | 5 (2.9%) |
| Other | 11 (6.4%) |
| **Graft type** |  |
| Unmodified bone marrow or PBSC | 86 (50%) |
| T-cell depleted PBSC | 72 (42%) |
| Cord blood | 13 (7.6%) |
| **Intensity of conditioning regimen** |  |
| Ablative | 99 (58%) |
| Reduced intensity | 51 (30%) |
| Nonmyeloablative | 21 (12%) |
| **GVHD prophylaxis** |  |
| CNI-based | 58 (34%) |
| PTCy-based | 41 (24%) |
| T-cell depleted PBSC | 72 (42%) |
| **Acute GVHD grade** |  |
| 0 | 67 (39%) |
| 1 | 28 (16%) |
| 2 | 63 (37%) |
| 3 | 11 (6.4%) |
| 4 | 2 (1.2%) |
| **Stage of Lower-GI acute GVHD** |  |
| 0 | 142 (83%) |
| 1 | 17 (9.9%) |
| 2 | 7 (4.1%) |
| 3 | 3 (1.8%) |
| 4 | 2 (1.2%) |

<sup>1</sup>Median (Q1, Q3); n (%)
CNI, calcineurin inhibitor; MDS/MPN, myelodysplastic/myeloproliferative neoplasms; PBSC, peripheral blood stem cell; PTCy, post-transplant cyclophosphamide.

We first examined the association between pre-transplant (day -7 to -1) dietary intake and clinical outcomes. To account for the skewed distribution and high fluctuation of daily nutrient intake, we calculated the average natural logarithm (ln) transformed pre-transplant intake of major macronutrients (i.e., carbohydrates, protein, fat, sugar, and fiber) using 826 patient-days of dietary intake (5.4 patient-days per patient). Multivariable regression models revealed significant associations between higher average pre-transplant intake of most macronutrients and improved OS, adjusted for the duration of pre-transplant broad-spectrum antibiotic exposure and clinical covariates (**Fig. 1b, Supplemental Table 1**). However, only fiber intake was significantly associated with improved OS (adjusted p=0.014, HR=0.469) and reduced incidence of severe acute GVHD (grade 2 or above, adjusted p=0.032, HR=0.480) for patients receiving non-T-cell depleted (non-TCD) grafts (**Table 2**; **Fig. 1c–d**). In contrast, we found no significant associations between post-transplant intake of the major macronutrients (day 0 to day 30) and the survival endpoints landmarked at day 30 when adjusted for the pre-transplant intake of the same macronutrient (**Supplemental Fig. 2, Supplemental Table 2**). These observations suggest the critical importance of nutritional intake during the pre-transplant period.

**Supplemental Figure 2.**
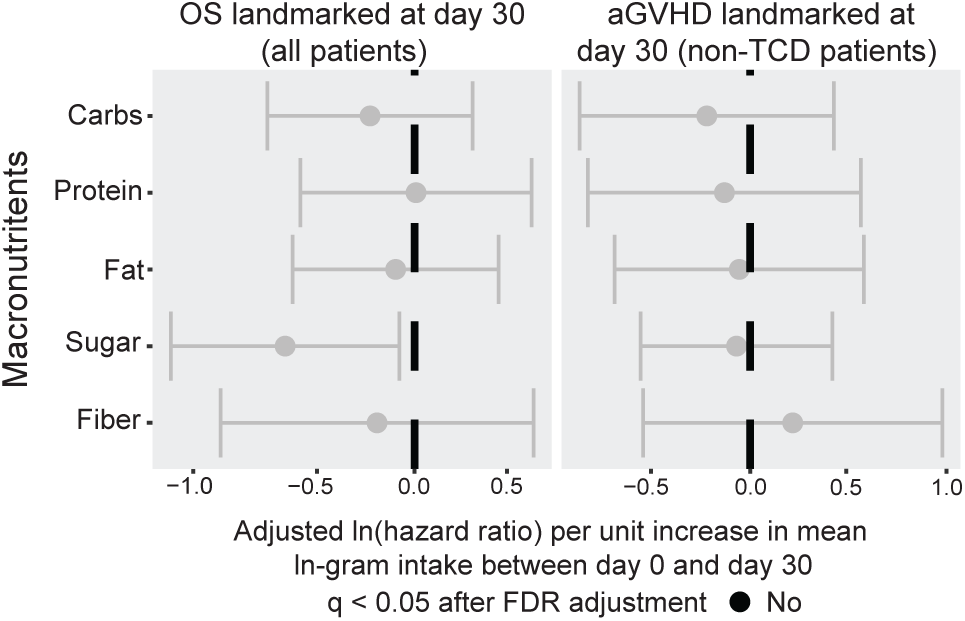
Post-transplant dietary patterns are not associated with OS or aGVHD. Associations between overall survival (OS, of all patients), severe acute GVHD (aGVHD, of non-TCD patients) and average ln-transformed post-transplant (days 0 to +30) daily intake (ln-grams/day) of macronutrients: carbohydrates, protein, fat, sugar, and fiber. Ln-transformed hazard ratios or sub-distributional hazard ratios) per unit increase in ln-gram intake were estimated for OS and aGVHD via multivariable Cox and Fine-Gray regression models, respectively. All models were adjusted for day -7 to day 30 broad-spectrum antibiotic exposure duration, pre-transplant dietary intake of the same macronutrient, disease type, conditioning intensity, and graft source (for OS models only). P-values were adjusted across macronutrients for each time-to-event outcome. No: Not significant.

**Table 2.**
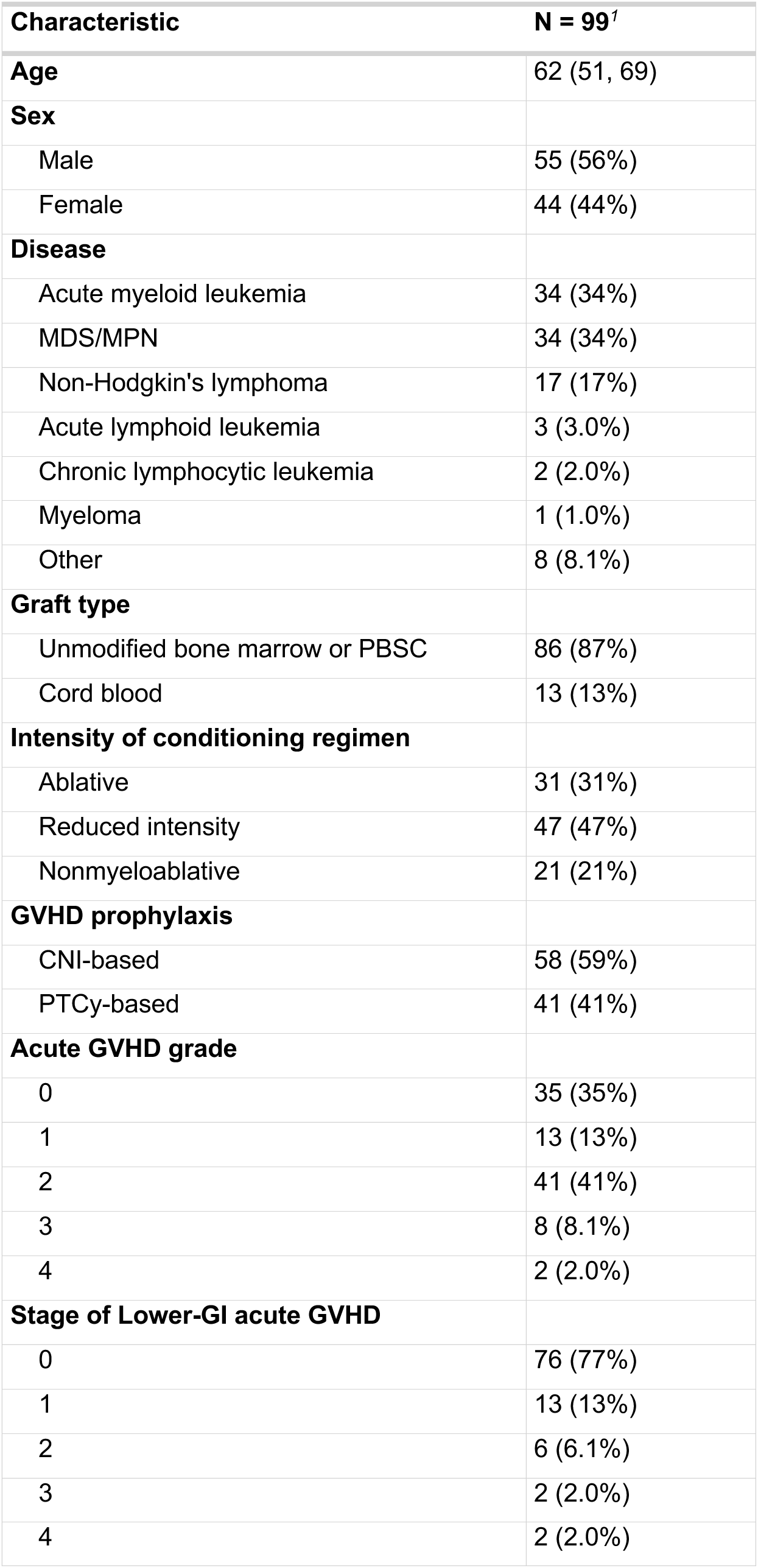
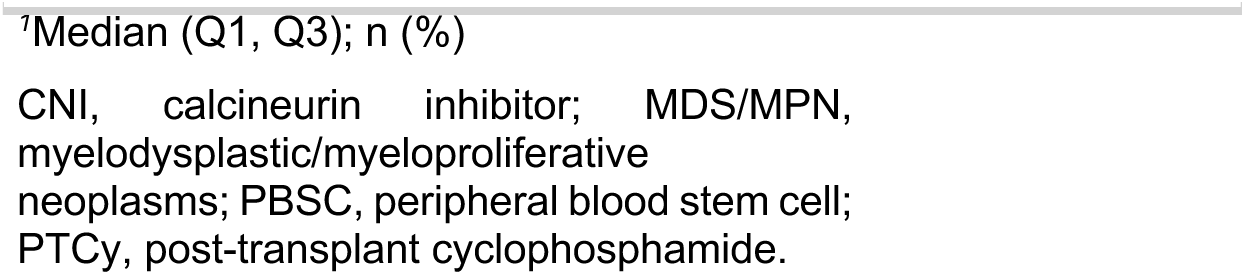
Characteristics of Non-TCD patients with available pre-HCT dietary data.

**Table 3.**
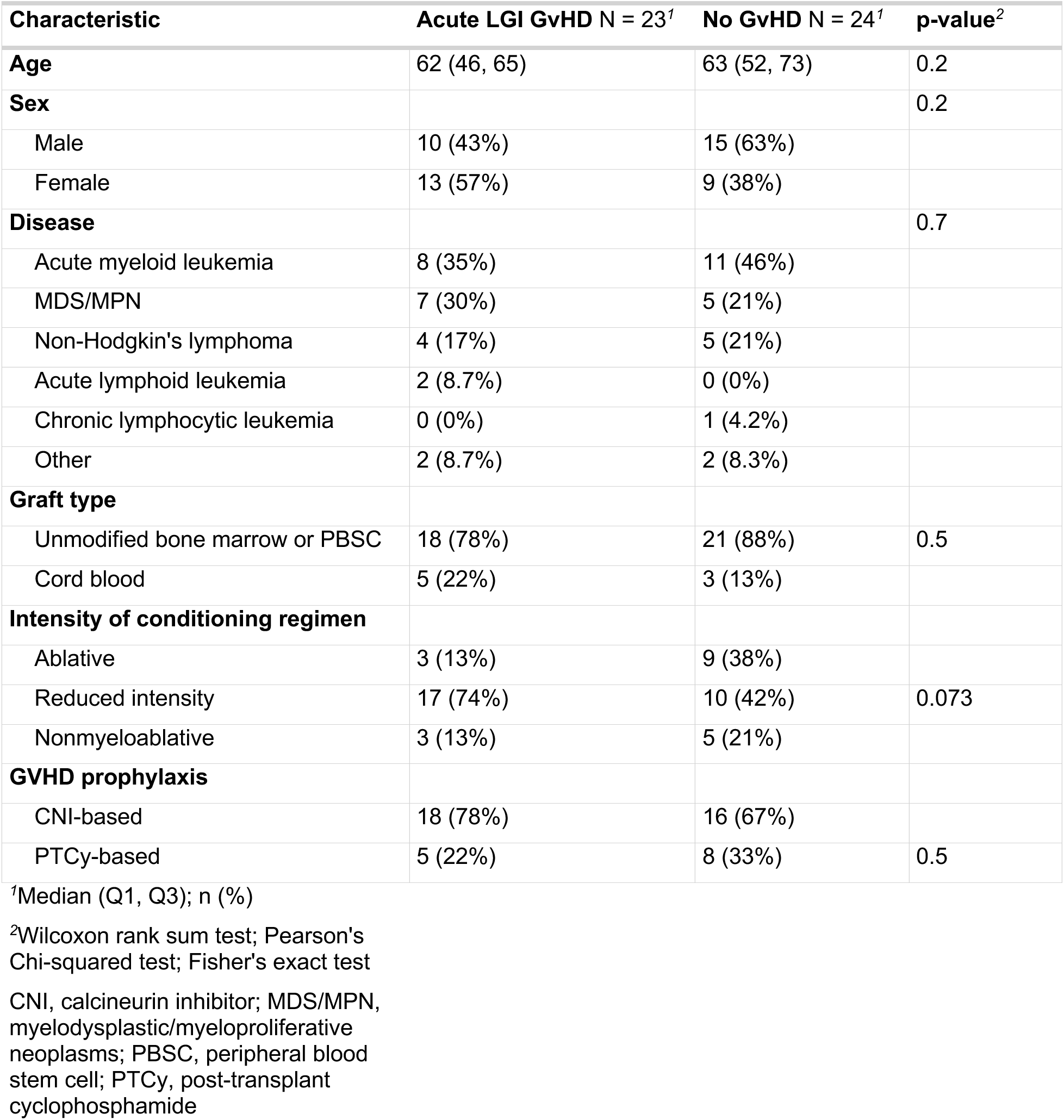
Characteristics of LGI-GVHD and Non-GVHD patients with available data from SCFA quantification.

| Characteristic | Acute LGI GvHD N = 23<sup>1</sup> | No GvHD N = 24<sup>1</sup> | p-value<sup>2</sup> |
| --- | --- | --- | --- |
| Age | 62 (46, 65) | 63 (52, 73) | 0.2 |
| Sex |  |  | 0.2 |
| Male | 10 (43%) | 15 (63%) |  |
| Female | 13 (57%) | 9 (38%) |  |
| Disease |  |  | 0.7 |
| Acute myeloid leukemia | 8 (35%) | 11 (46%) |  |
| MDS/MPN | 7 (30%) | 5 (21%) |  |
| Non-Hodgkin's lymphoma | 4 (17%) | 5 (21%) |  |
| Acute lymphoid leukemia | 2 (8.7%) | 0 (0%) |  |
| Chronic lymphocytic leukemia | 0 (0%) | 1 (4.2%) |  |
| Other | 2 (8.7%) | 2 (8.3%) |  |
| Graft type |  |  |  |
| Unmodified bone marrow or PBSC | 18 (78%) | 21 (88%) | 0.5 |
| Cord blood | 5 (22%) | 3 (13%) |  |
| Intensity of conditioning regimen |  |  |  |
| Ablative | 3 (13%) | 9 (38%) |  |
| Reduced intensity | 17 (74%) | 10 (42%) | 0.073 |
| Nonmyeloablative | 3 (13%) | 5 (21%) |  |
| GVHD prophylaxis |  |  |  |
| CNI-based | 18 (78%) | 16 (67%) |  |
| PTCy-based | 5 (22%) | 8 (33%) | 0.5 |

<sup>1</sup>Median (Q1, Q3); n (%)
<sup>2</sup>Wilcoxon rank sum test; Pearson's Chi-squared test; Fisher's exact test
CNI, calcineurin inhibitor; MDS/MPN, myelodysplastic/myeloproliferative neoplasms; PBSC, peripheral blood stem cell; PTCy, post-transplant cyclophosphamide

### Cellulose consumption is positively associated with enrichment of beneficial commensal bacteria and fecal concentrations of SCFAs

Given the dynamic microbiome shifts that occur during hospitalization after transplantation^25,27,30,31,32,33,34,35,36^, we hypothesized that pre-transplant fiber consumption could improve microbial diversity, expansion of butyrate-producing commensals, and suppression of pathogenic taxa, which are microbial features previously associated with improved transplant outcomes^27,33,37^. We therefore analyzed the microbial dynamics of longitudinal fecal samples collected between days 0 and 30, the early post-transplant phase during which patients are highly susceptible to microbial dysbiosis, infections, and GVHD ^27,33,38,39,40^.

We evaluated the correlation between pre-transplant (day -7 to 0) average fiber consumption and intestinal microbial α-diversity of 921 longitudinal stool samples in the early post-transplant period via generalized estimating equations (GEEs) while adjusting for antibiotic exposure and time of fecal sample collection. In addition to overall fiber intake, we considered soluble and insoluble fiber subgroups to further explore associations with microbial diversity. Higher average pre-transplant intake of total fiber (p=0.135) or of soluble fiber (p=0.478) did not correlate with increased microbial α-diversity of samples collected on post-transplant days 0 and 30 (**Fig. 2a**). However, higher average pre-transplant intake of insoluble fiber correlated with increased α-diversity post-transplant (p=0.024), accompanied by reduced relative abundance of *Enterococcus* (p=0.032, **Fig. 2b**) and greater relative abundance of *Blautia* (p=0.004, **Fig. 2b**). These results suggest that insoluble fiber intake during the pre-transplant period may modulate the post-transplant taxonomical composition of the gut microbiome. We found no association between pre-transplant intake of protein, fat, sugar, or other carbohydrates and the post-transplant trajectory of α-diversity or the relative abundances of *Enterococcus* or *Blautia* between days 0 and 30 (**Supplemental Fig. 3a**).

**Figure 2.**
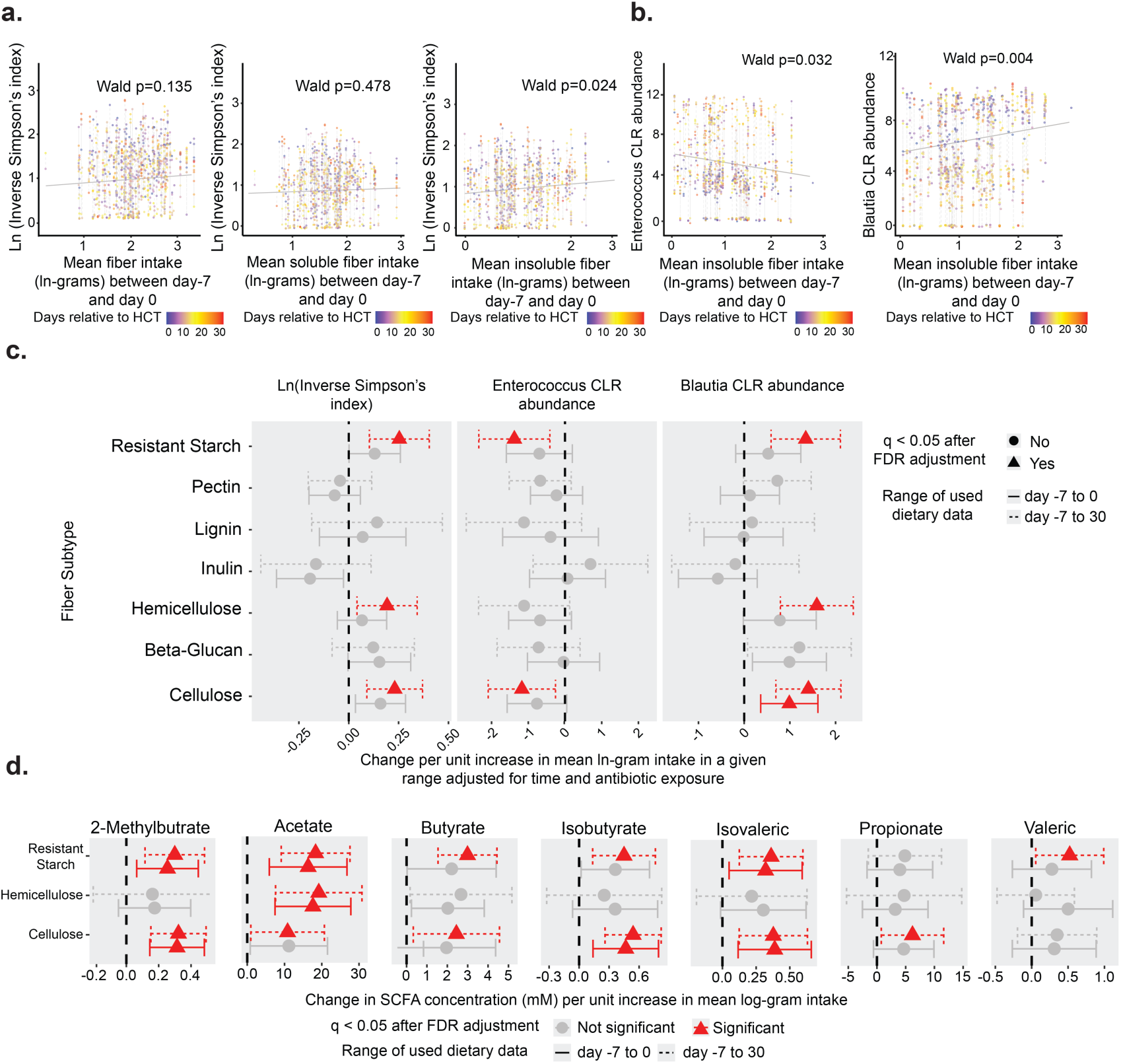
Intake of insoluble fiber and cellulose is associated with increased microbial diversity, enrichment of beneficial commensal bacteria, suppression of pathobionts, and enhanced production of SCFAs during allo-HCT. a: Associations between average ln-transformed pre-HCT (days -7 to 0 relative to HCT) fiber, soluble fiber, and insoluble fiber intake (ln-grams/day) and post-HCT (days 0 to 30) α-diversity (inverse Simpson’s index). (N=923 samples from 159 patients with available pre-HCT dietary data and post-transplant microbiome data.) b: Associations between average ln-transformed pre-HCT (days -7 to 0 relative to HCT) insoluble fiber intake (ln-grams/day) and post-HCT (days 0 to 30) centered log-ratio (CLR) transformed *Enteroccocus* and *Blautia* abundances (N=923 samples from 159 patients). c: Associations between day 0 to day 30 longitudinal microbial features (inverse Simpson’s index, CLR-transformed *Enterococcus* and *Blautia* abundances) and average intake of 7 major fiber subtypes during the pre-HCT window (days -7 to 0, solid line, N=923 samples from 159 patients) and during the extended peri-transplant window (days -7 to 30, dashed line, N=935 samples from 161 patients with available peri-transplant dietary data and post-transplant microbiome data). P-values were adjusted across fiber subtypes for each microbial feature outcome. Fiber subtypes for which FDR-adjusted p-values < 0.05 are shown with estimated effect sizes annotated by red triangles with red unadjusted 95% confidence intervals. d: Associations between day 0 to day 30 longitudinal short-chain fatty acid (SCFA) concentrations and average intake of 3 fiber subtypes during the pre-HCT window (days -7 to 0, solid line, N=202 GC-MS samples from 45 patients with available pre-HCT dietary data and post-transplant SCFA data) and during the extended peri-transplant window (days -7 to 30, dashed line, N=204 GC-MS samples from 46 patients with available peri-transplant dietary data and post-transplant SCFA data). P-values were adjusted across fiber subtypes for each SCFA. Fiber subtypes for which FDR-adjusted p-values < 0.05 are shown with estimated effect sizes annotated by red triangles with red unadjusted 95% confidence intervals. All microbial and SCFA features were quantified by 16S rRNA sequencing and GC-MS of fecal samples collected longitudinally between day 0 to 30, respectively. Generalized estimating equations (GEE) of longitudinal microbial and SCFA features were adjusted for the duration of broad-spectrum antibiotic exposure from day -7 to day 30 and sample collection days relative to HCT. For panels a and b, each point in the scatter plot represents a fecal sample, colored by days of collection relative to HCT. Samples collected from the same patient are connected by dashed lines. The solid, grey-colored line represents the expected value of microbial features at each average intake value based on the GEE models, with sample collection time fixed at the midpoint of the time window (day 15) and antibiotic exposure fixed at the median duration of 12 days of exposure between day –7 to day 30 relative to HCT.

**Supplemental Figure 3.**
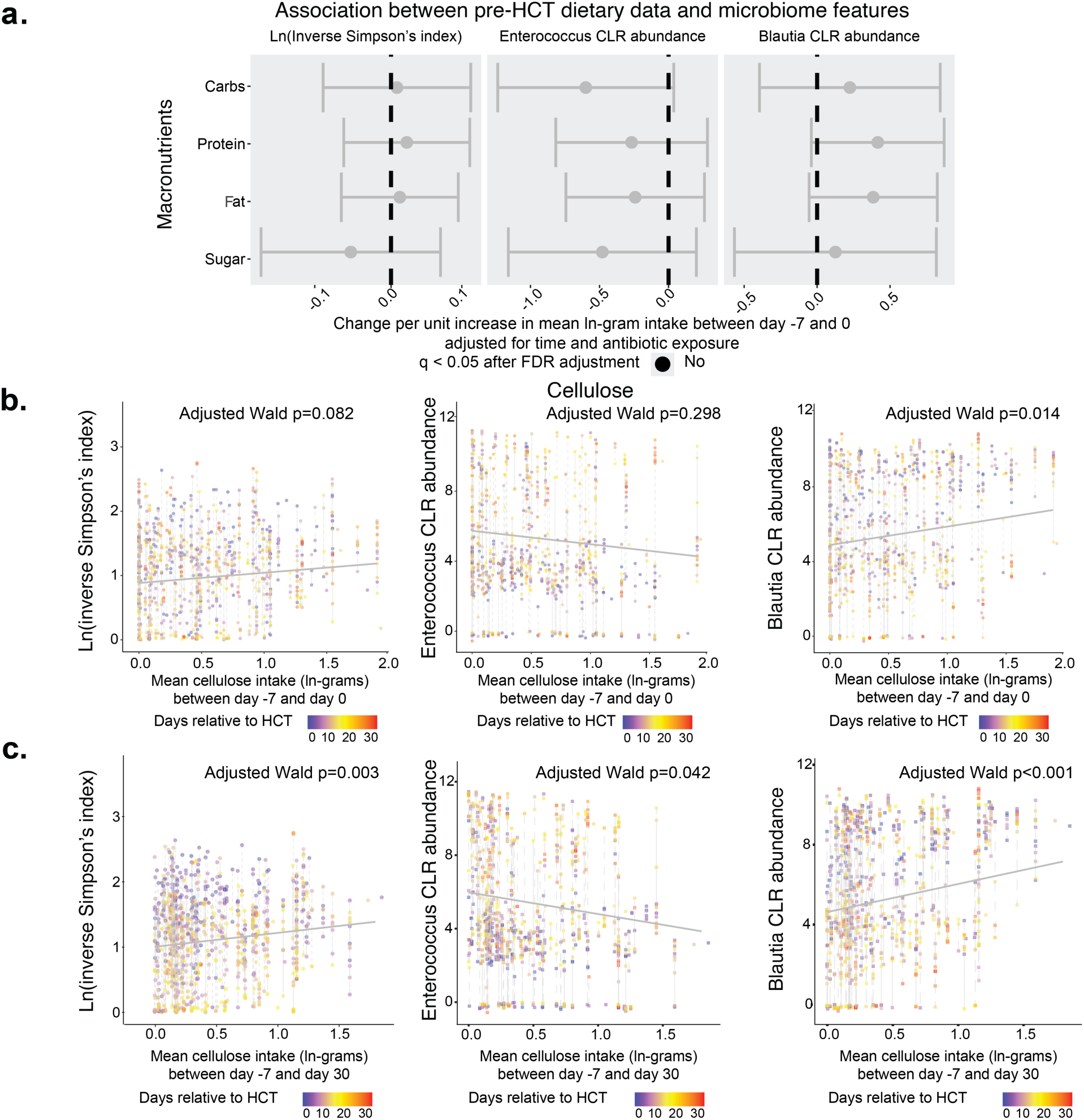
Cellulose intake is associated with increased microbial diversity, enrichment of beneficial commensals, and suppression of pathobionts pre- transplantation. a: Associations between day 0 to day 30 longitudinal microbial features (inverse Simpson’s index, CLR-transformed *Enterococcus* and *Blautia* abundances) and average intake of 4 macronutrients during the pre-HCT window (days -7 to 0). P-values were adjusted across macronutrients for each microbial feature outcome. (N=923 samples from 159 patients with available pre-HCT dietary data and post-transplant microbiome data.) No: not significant. b: Associations between average ln-transformed pre-HCT (days -7 to 0) cellulose intake (ln-grams/day) and α-diversity (inverse Simpson’s index), centered log-ratio (CLR) transformed *Enteroccocus* and *Blautia* abundances (N=923 samples from 159 patients). c: Associations between average ln-transformed full follow-up (days -7 to 30) cellulose intake (ln-grams/day) and α-diversity (inverse Simpson’s index), centered log-ratio (CLR) transformed *Enteroccocus* and *Blautia* abundances. (N=935 samples from 161 patients with available peri-transplant dietary data and post-transplant microbiome data.) All microbial and SCFA features were quantified by 16S rRNA sequencing and GC-MS of fecal samples collected longitudinally between day 0 to 30, respectively. Generalized estimating equations (GEEs) of longitudinal microbial and SCFA features were adjusted for the duration of broad-spectrum antibiotic exposure from day -7 to day 30 and sample collection days relative to HCT. In b and c, each point in the scatter plot represents a fecal sample, colored by days of collection relative to HCT. Samples collected from the same patient were connected by dashed lines. The solid, grey-colored line represents the expected value of microbial features at each average intake value based on the GEE models, with sample collection time fixed at the midpoint of the time window (day 15) and antibiotic exposure fixed at the median duration of 12 days of exposure between day -7 to day 30 relative to HCT.

Next, we used GEEs to further evaluate the associations between microbial dynamics (e.g., α-diversity, *Blautia* abundance) and the most relevant fiber subtypes, including resistant starch, pectin, lignin, inulin, hemicellulose, beta-glucan, and cellulose^41^. We calculated the average daily intake during the pre-transplant period (days -7 to -1) (**Supplementary Table 1**) and the extended-transplant window (days -7 to 30, **Supplementary Table 2**). Among the subtypes, only cellulose, resistant starch, and hemicellulose (classified as insoluble or mixed fibers) exhibited significant associations with microbial features (**Fig. 2c**). Notably, we found a significant positive association between pre-transplant intake of cellulose, a fully insoluble fiber and major component of plant-based diets known to enrich for anaerobic and SCFA-producing taxa^42^, and *Blautia* abundance (adjusted p=0.014, **Supplemental Fig. 3b**). Cellulose was the only fiber subtype for which we found any significant association between the average pre-transplant intake and any aspect of microbial dynamics. Moreover, higher average intake of cellulose between days -7 and 30 was significantly associated with higher α-diversity (adjusted p=0.003), lower *Enterococcus* abundance (adjusted p=0.042), and higher *Blautia* abundance (adjusted p<0.001) (Supplemental Fig. 3c).

We also investigated the association of cellulose, resistant starch, and hemicellulose with production of SCFAs, which are microbial-derived metabolites that result from fiber fermentation and have protective effects within the intestine^43,44^. Using gas chromatography-mass spectrometry (GC-MS), we quantified fecal concentrations of acetate, butyrate, propionate, 2-methylbutyrate, isobutyrate, and isovaleric and valeric acids in samples collected from day 0 to 30 relative to transplantation from non-TCD patients (204 samples from 46 patients). We found significant associations between elevated levels of all measured SCFAs and intake of at least one of the three selected fiber subtypes, either during the pre-transplant or peri-transplant phase (**Fig. 2d**). Notably, cellulose intake was positively associated (p<0.05) with all but one of the tested SCFAs (**Supplemental Fig. 4**). In summary, we demonstrated that higher than average pre-transplant intake of insoluble fiber, particularly cellulose, was associated with increased microbial diversity, reduced abundance of *Enterococcus*, and enrichment of *Blautia* during the early post-transplant period in allo-HCT recipients. In addition, cellulose intake across the allo-HCT timeline correlated with higher concentrations of SCFAs.

**Supplemental Figure 4.**
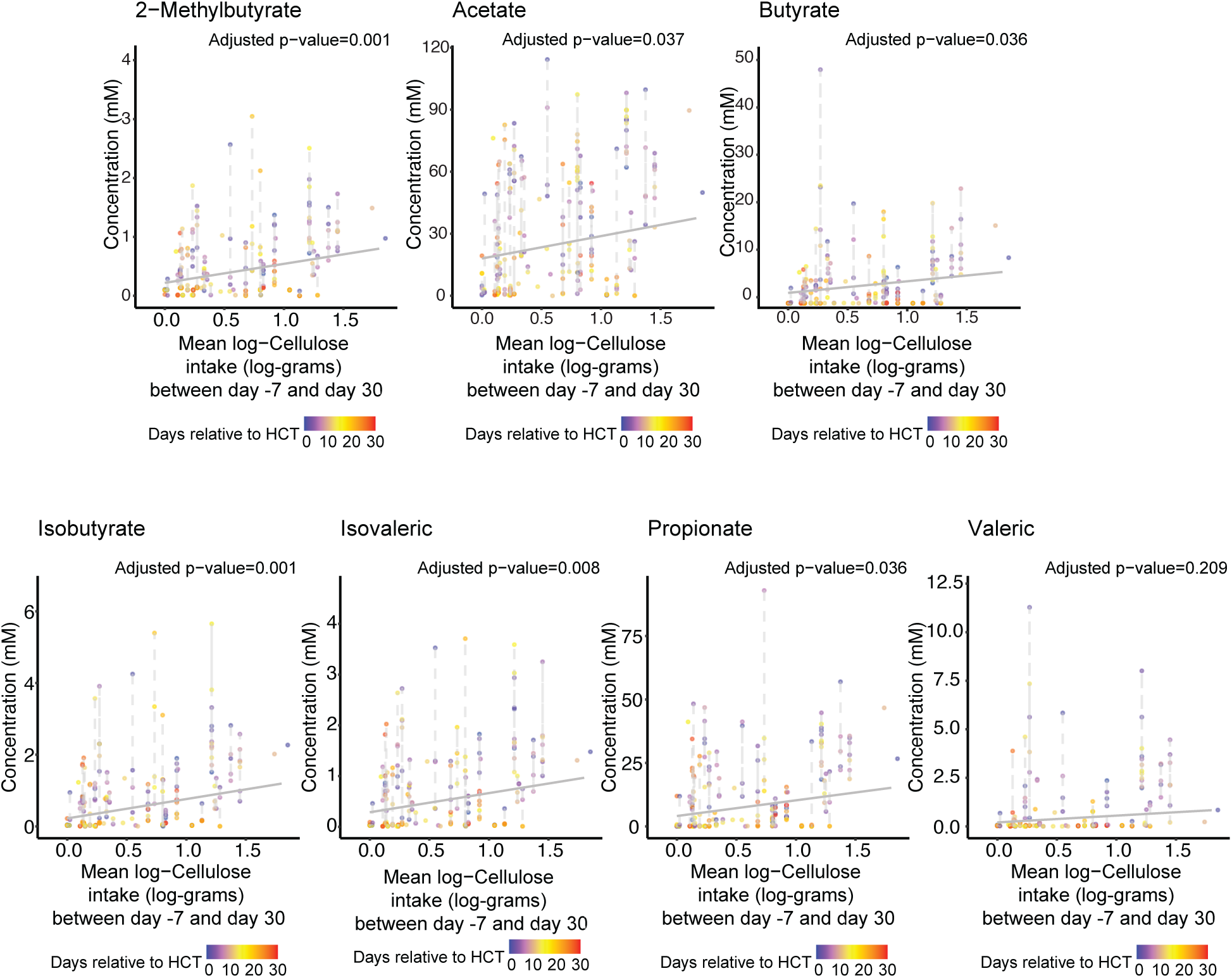
Cellulose intake is positively associated with increased concentrations of fecal SCFAs in non-T cell depleted allo-HCT patients. Associations between average ln-transformed cellulose intake (grams/day) between days -7 to 30 relative to HCT and fecal concentrations of SCFAs. SCFA concentrations were quantified from fecal samples collected longitudinally between day 0 to 30. GEEs of SCFA concentrations were adjusted for the duration of broad-spectrum antibiotic exposure from day -7 to day 30 and sample collection days relative to HCT. Each point in the scatter plot represents a fecal sample, colored by days of collection relative to HCT. Samples collected from the same patient are connected by dashed lines. The solid, grey-colored line represents the expected value of SCFA concentration at each average intake value based on the GEE models, with sample collection time fixed at the midpoint of the time window (day 15) and antibiotic exposure fixed at the median duration of 12 days of exposure between day -7 to day 30 relative to HCT. GC-MS in samples collected from day 0 to 30 relative to transplantation from non-TCD patients (N=204 GC-MS samples from 46 patients with available peri-transplant dietary data and post-transplant SCFA data).

### SCFA fecal concentrations are depleted in patients with LGI-GVHD

Given our observed association between cellulose intake and enhanced SCFA production in non-TCD allo-HCT recipients, and the localization of SCFA production to the colon, where fiber fermentation occurs^28^, we hypothesized that patients with LGI-GVHD would exhibit reduced levels of these key microbial metabolites. To test this, we analyzed 241 fecal samples from non-TCD allo-HCT recipients collected between days -7 to 30 and compared the fecal SCFA concentrations of patients who developed LGI-GVHD (n=23) or did not develop acute GVHD (n=24). Patients with LGI-GVHD exhibited lower concentrations of acetate, butyrate, valeric acid propionate, isobutyrate, isovaleric, and 2-methylbutyrate; the reduced levels of acetate, butyrate, and valeric acid were statistically significant (adjusted p<0.05) (**Fig. 3**, **Supplemental Fig. 5**).

**Figure 3.**
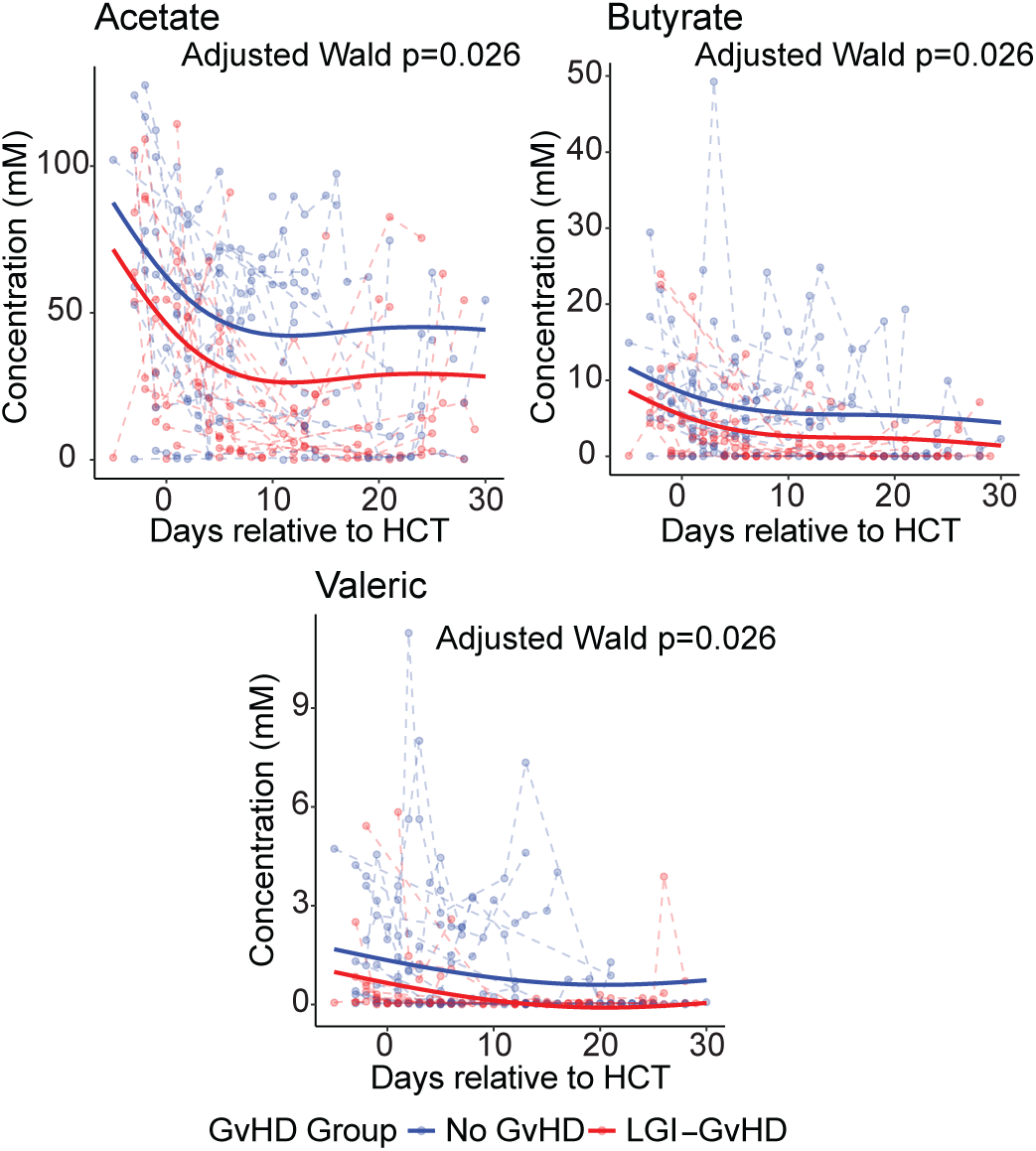
Patients with lower gastrointestinal GVHD (LGI-GVHD) exhibit significantly reduced fecal concentrations of SCFAs acetate, butyrate and valeric after allo-HCT. Longitudinal trajectories of fecal acetate, butyrate, and valeric acid in millimolar (mM) concentration between day -7 and 30 relative to transplant (N=241 samples from 47 patients; blue, no acute GVHD [n=24]; red, LGI-GVHD [n=23]). Dots represent stool samples. Each dashed line represents the trajectory of a single patient. Solid blue and red curves represent the mean trajectories of SCFA concentration for the two patient groups based on generalized estimating equation (GEE) models, assuming a constant group-wise difference. The Wald p-values associated with the difference between No acute GVHD and LGI-GVHD groups were adjusted across the 7 measured SCFAs. Analysis was restricted to allo-HCT recipients who did not receive T cell–depleted (non-TCD) grafts (n=101), enabling assessment of GVHD- associated microbiome and metabolite changes without confounding from graft manipulation.

**Supplemental Figure 5.**
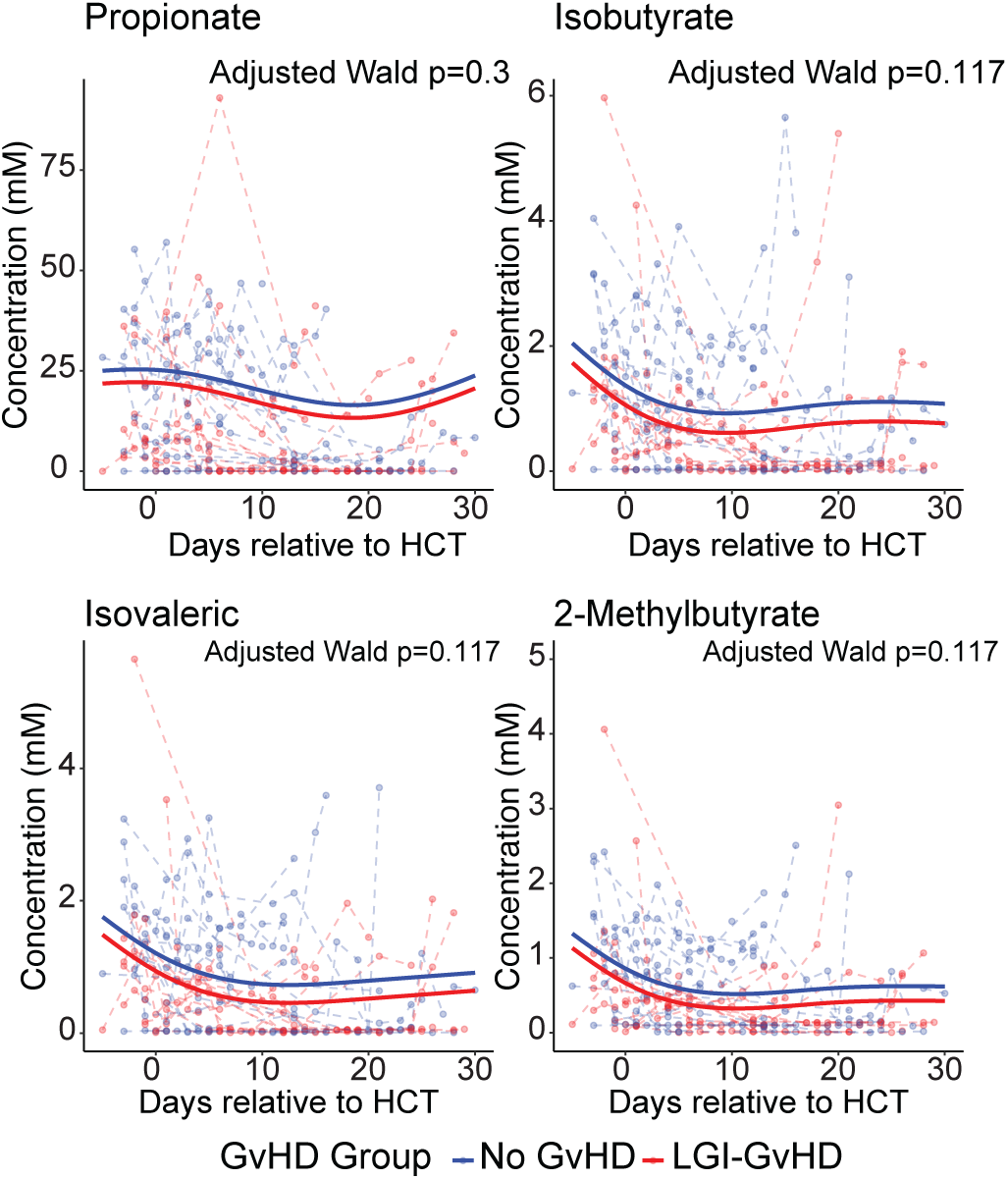
Patients with lower gastrointestinal GVHD (LGI-GVHD) have lower concentrations, while not significant, of fecal SCFA after allo-HCT. Longitudinal trajectories of fecal propionate, isobutyrate, isovaleric acid, and 2-methylbutyrate in millimolar (mM) concentrations between day -7 and 30 relative to transplant (N=241 samples from 47 patients; blue, no acute GVHD [n=24]; red, LGI-GVHD [n=23). Dots represent stool samples. Each dashed line represents the trajectory of a single patient. Solid blue and red curves represent the mean trajectories of SCFA concentration for the two patient groups based on generalized estimating equation (GEE) models assuming a constant group-wise difference. The Wald p-values associated with the difference between No GVHD and LGI-GVHD groups were adjusted across the 7 measured SCFAs.

Collectively, in our clinical allo-HCT cohort, we observed positive associations between cellulose intake and increased microbial diversity, reduced abundance of *Enterococcus*, and enhanced SCFA production. In addition, patients with LGI-GVHD had reduced levels of beneficial microbial metabolites.

### A 12% cellulose diet reproduces human microbiome signatures and improves GVHD survival in pre-clinical models

To further our mechanistic understanding of how the intestinal microbiome changes in response to dietary fiber in allo-HCT, we next evaluated the impact of cellulose supplementation in a preclinical mouse model of allo-HCT. We used an MHC-disparate mouse model of allo-HCT (C57BL/6J donors into BALB/c recipients), with mice receiving bone marrow only (BM) or bone marrow plus T cells (BM+T) to induce GVHD. We tested cellulose alone as the dietary fiber source, because we had observed the strongest association between cellulose and clinical outcomes in allo-HCT patients. Cellulose is the most common type of fiber^45,46^. It cannot be digested by mammalian enzymes, and its digestion relies entirely on microbial fermentation in the gut^42^. Mice were fed customized diets 2 days pre-transplant (conditioning period) containing 0% (fiber-free), 6% (comparable to standard chow), or 12% (high-fiber) cellulose while maintaining similar concentrations of other macronutrients (**Fig. 4a**). Mice were followed for 90 days post-transplant for GVHD morbidity and mortality. The BM+T 12% cellulose group exhibited significantly improved survival compared to BM+T mice receiving 0% (p=0.016) or 6% (p=0.022) cellulose (**Fig. 4b**). There was no difference in survival between the 0% and 6% groups, indicating that the standard dietary fiber concentration was insufficient to confer a protective effect.

**Figure 4.**
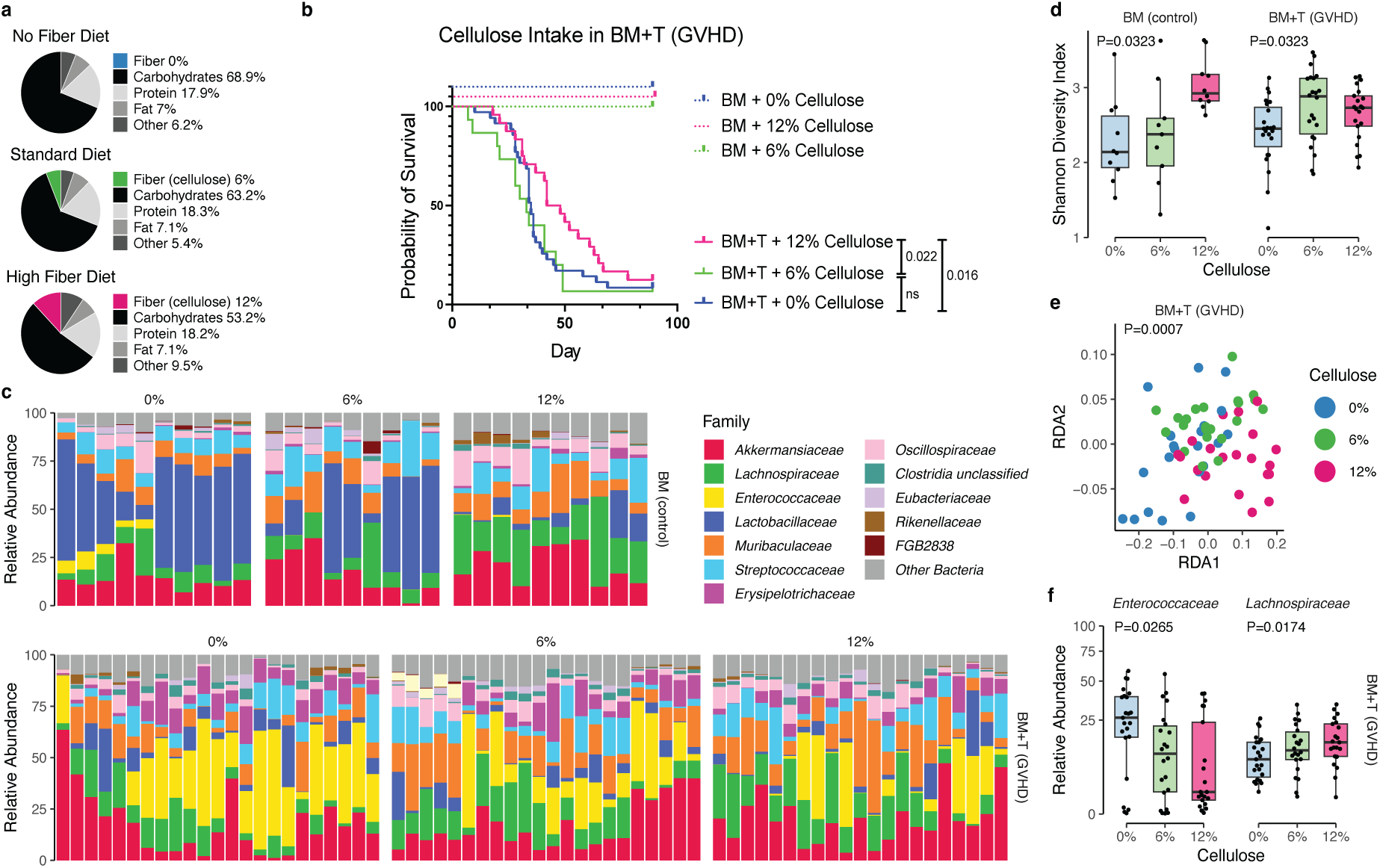
A 12% cellulose diet mirrors human allo-HCT results by improving GVHD survival, enhancing microbial diversity, and enriching for beneficial taxa. a: Composition of experimental diets containing 0%, 6%, or 12% cellulose. Macronutrient content (fat, protein, non-fiber carbohydrates) was held similar across diets to isolate the effect of fiber concentration. b: Mice were assigned to experimental diets and monitored for survival for 90 days post-transplant. Cohorts included BM only (n=15 per group) and BM+T (n=30 per group), total N=135 from 3 independent experiments. Logrank (Mantel-Cox) test. c: Microbiome composition at day +7 post-transplant based on shotgun metagenomic sequencing of fecal samples (n=95) from BM mice (0% n=9, 6% n=9, 12% n= 10) and BM+T mice (0% n=23, 6% n=22, 12% n=21). |d: Microbial α-diversity (Shannon index) comparisons between dietary groups at day +7 post-BMT. Benjamini-Hochberg adjusted p-values computed for cellulose coefficients in generalized linear models. e: Redundancy analysis (RDA) of microbial community structure using a reduced model. Significance evaluated by permutation-based ANOVA. f: Relative abundance of specific bacterial families at day +7 post-BMT in GVHD mice. Benjamini-Hochberg adjusted p-values computed for cellulose coefficients in generalized linear models.

To assess whether mouse microbiome changes mirrored those seen in allo-HCT patients, we performed shotgun metagenomic sequencing on fecal samples collected at day 7 post-transplant, a time point associated with the initiation of early GVHD pathogenesis (**Fig. 4c**). Microbial composition diverged significantly across diet groups, independent of transplant condition (BM or BM+T). Microbial α-diversity increased with cellulose concentration in both the BM-only control (p=0.0323) and BM+T (p=0.0323) mice (**Fig. 4d**). In addition, intestinal microbiome taxonomical distributions in GVHD mice at day 7 post-transplant significantly differed with cellulose concentration (p=0.0007) (**Fig. 4e**). These diversity gains were accompanied by shifts in microbial taxa consistent with our human results, including enriched abundance of *Lachnospiraceae*, the family that includes *Blautia* (p=0.0174), and reduced abundance of *Enterococcaceae*, the family including *Enterococcus* (p=0.0265), in a cellulose concentration-dependent manner (**Fig. 4f**). Thus, these murine experiments recapitulated the observations from our allo-HCT patient cohort, demonstrating that a higher fiber diet (12% cellulose) enhances survival in the context of GVHD and promotes colonization by beneficial taxa (*Lachnospiraceae)* while suppressing colonization by pathogenic species (*Enterococcaceae)*.

### A 12% cellulose diet promotes butyrate production and epithelial integrity in the large intestine

We next investigated whether the taxonomic changes corresponded to shifts in the metabolic potential of the microbiota. Using the regularized log-ratio regression method FLORAL^47^, we modeled the percentage of cellulose intake and conducted feature selection for the pathways detected from the shotgun metagenomic data. We found that increasing dietary cellulose enriched metabolic pathways involved in polysaccharide breakdown and beneficial metabolite production in murine intestinal microbial communities (**Fig. 5a**). Among the positively associated pathways were hexuronide and hexuronate degradation, which enable microbes to catabolize plant-derived complex carbohydrates into fermentable monosaccharides. Also enriched was the biosynthesis of thiamine diphosphate (vitamin B1), an essential microbial cofactor involved in energy metabolism and host–microbe symbiosis. Notably, the pathway for *Clostridium acetobutylicum* acidogenic fermentation, which supports SCFA production including butyrate, was also associated with cellulose intake.

**Figure 5.**
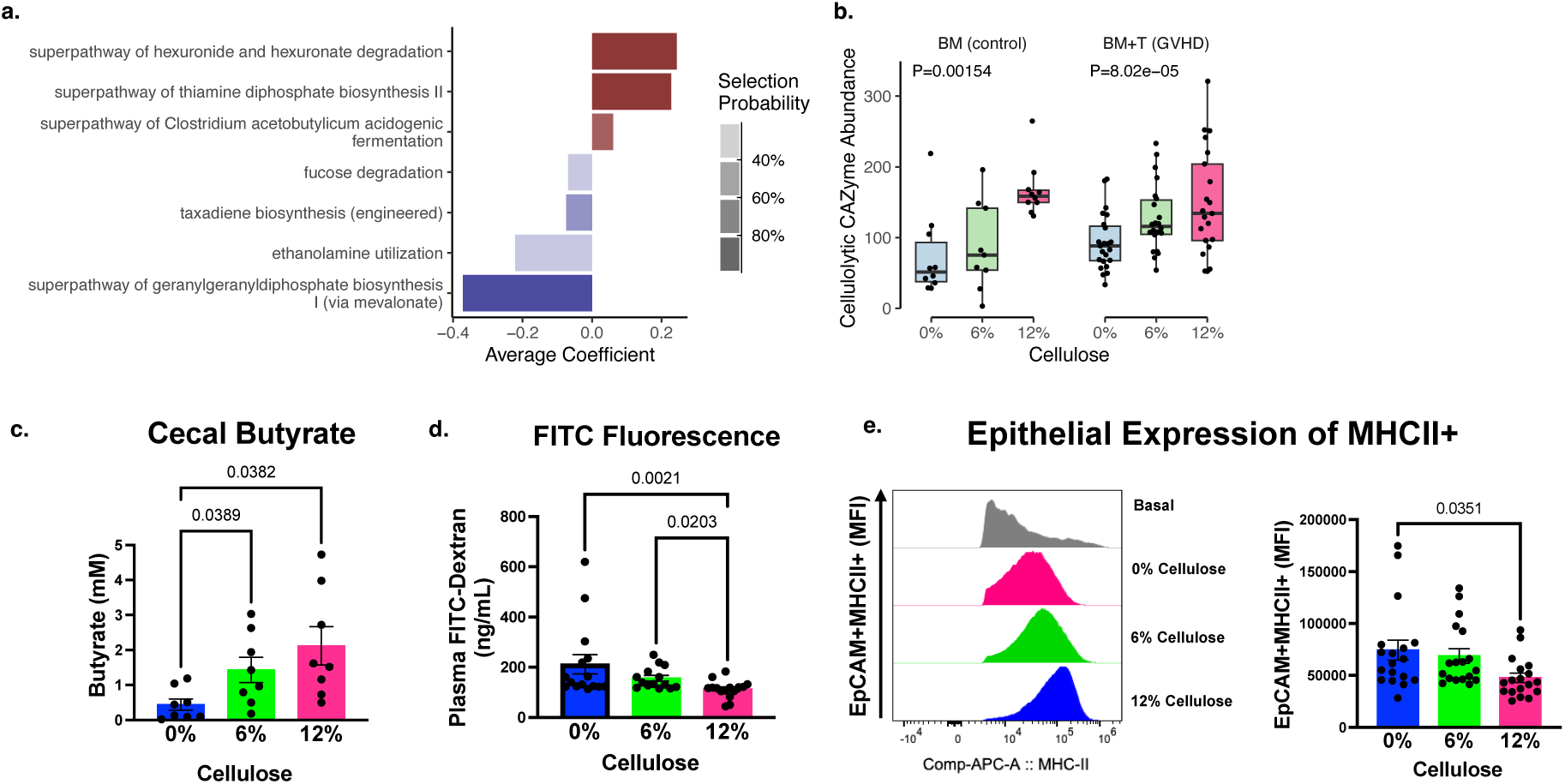
A 12% cellulose diet promotes epithelial cell homeostasis and intestinal barrier integrity in the large intestine of GVHD mice. a: Pathway selection probabilities and average coefficients from cross-fold validation of FLORAL models of pathway abundance *vs.* cellulose percentage. b: Cellulolytic CAZyme gene abundance across diet compositions. Benjamini-Hochberg adjusted p-values computed for cellulose coefficients in generalized linear models. c: Quantification of cecal butyrate concentrations. Cecal samples were collected at day 7 post allo-HCT. GC-MS analysis, N=32 (n=8 per group), from 3 independent experiments. One-Way ANOVA followed by Tukey’s test. d: FITC-Dextran assay measuring epithelial permeability by plasma fluorescence levels. N=45 (n=15 per group), from 3 independent experiments. One-Way ANOVA followed by Tukey’s test. e: Flow cytometry quantification of inflammatory epithelial cells (EpCAM ⁺CD45⁻MHCII⁺) of GVHD mice, day 7 post BMT, treated with 0%, 6% and 12% cellulose. Representative histogram overlays of MHC-II staining across dietary groups. Non-transplanted mice fed standard diet, were included for reference (basal expression). N=45 (n=15 per group), from 3 independent experiments. One-Way ANOVA followed by Tukey’s test.

Conversely, pathways negatively associated with cellulose intake included fucose degradation and ethanolamine use, both of which are frequently exploited by facultative pathogens during gut inflammation^48,49^. Fucose degradation can reduce protective mucus layers in the intestine, while ethanolamine use provides a competitive advantage for enteric pathogens in dysbiotic environments^50,51^. We also observed a reduction in the taxadiene biosynthesis and geranylgeranyl diphosphate biosynthesis pathways, which are involved in microbial isoprenoid metabolism and have been linked to bacterial virulence and host immune modulation^52,53,54^.

To link these pathway shifts to microbial capacity for cellulose metabolism, we used dbCAN3^55^ to annotate carbohydrate-active enzymes (CAZymes) and quantify the abundance of microbial genes encoding enzymes that degrade cellulose. Microbiota from mice fed cellulose diets had significantly higher abundance of genes encoding cellulolytic enzymes in a cellulose concentration dependent manner in the BM (p=0.00154) and in the BM+T groups (p<0.0001) (**Fig. 5b**), corroborating the observed functional shifts as a direct consequence of the dietary intervention. Subsequently, we tested the correlation between cellulose consumption and microbiome-derived metabolites, focusing on butyrate concentrations because they were lower in fecal samples from LGI-GVHD patients. Mice fed the 12% cellulose diet exhibited significantly higher levels of cecal butyrate compared to the 0% cellulose cohort (p=0.0382, **Fig. 5c**), further supporting the model in which dietary fiber promotes gut health via microbial fermentation and SCFA production.

We next focused on the biological effects of dietary cellulose on the LGI tract of GVHD mice. We hypothesized that a 12% cellulose diet would lead to improved epithelial health and barrier integrity, two key pathophysiological features of LGI-GVHD that are closely linked to microbiome injury and severity of inflammation^56^. Using a fluorescein isothiocyanate (FITC)-dextran assay to directly measure intestinal permeability, we found that plasma FITC levels 7 days after transplantation were significantly lower in mice fed 12% cellulose compared to those fed 0% (p=0.0021) and 6% cellulose (p=0.0203), indicating and enhanced barrier integrity (**Fig. 5d**).

We used flow cytometry to characterize epithelial inflammation by quantifying the intensity of MHC-II expression in epithelial cells (CD45⁻ EpCAM^+^) 7 days after transplant, which has been positively associated with GVHD severity^57,58^. Mice fed the 12% cellulose diet had a lower mean fluorescence intensity (MFI) of these epithelial cells compared to 0% controls (p=0.0351, **Fig. 5e**), while mice fed 0% versus 6% cellulose had no significant difference in MFI. Thus, we concluded that 12% cellulose preserves intestinal epithelial homeostasis and integrity and increases microbial butyrate production.

### A 12% cellulose diet suppresses inflammatory processes in GVHD mice

We next focused on the biological effects of dietary cellulose on inflammation in the LGI tract of GVHD mice. We performed single-cell RNA sequencing (scRNA-seq) of allogeneic T cells from large intestine tissues collected on day +7 post-transplant from GVHD mice fed diets containing 0%, 6%, or 12% cellulose (**Supplemental Fig. 6a**). We used clustering analysis to identify donor-derived (H-2Kb^+^) CD4^+^ and CD8^+^ T cells and Ki67^+^ (*Mki67*) to measure proliferation (**Supplemental Fig. 6b**). Notably, mice fed the 12% cellulose diet displayed a lower proportion of proliferating (Ki67^+^) allogeneic CD4^+^ and CD8^+^ T cells compared to mice fed 0% or 6% cellulose (**Supplemental Fig. 6c**). We further used the scRNA sequencing data to assess how dietary cellulose influences CD4^+^ T cell gene expression in the LGI tract as they are the main drivers of GVHD in our allo-HCT model^21,58,59,60,61,62,63^ (**Supplemental Fig. 6d**). Mice fed 12% cellulose, compared to 0% and 6%, had CD4^+^ T cells with significantly higher expression of genes associated with regulation of T cell activation (*Nr4a1*, [nuclear receptor 4A1] and *Nr4a3* [nuclear receptor 4A3]); transcription factors downstream of TCR activation that can suppress the expression of genes involved in cytokine (IFN-γ, IL-2) production; and genes that promote the development of T regulatory (Treg) cells^64,65^ (**Fig. 6a**). We found significant upregulation of *Maf* (MAF bZIP Transcription Factor), which is involved in T helper (Th) cell differentiation and cytokine regulation (e.g., positive regulator of IL-10 and a negative regulator of IL-2)^66^; *Bcl2* (B-cell lymphoma 2) and *Bcl2a1d* (Bcl2-related protein A1), which are cell apoptosis and cycle quiescence genes^67^; *Irf2bp2* (interferon regulatory factor 2 binding protein 2), a transcription cofactor involved in the regulation of T cell activation^68,69^; *Zfp36l1*, *Zfp36l2*, and *Zfp36*, which are tristetraprolin family members essential in Treg cells for immune homeostasis and autoimmunity prevention^70^; and *St8sia4*^71,72^ (polysialyltransferase IV). These findings suggest that a 12% cellulose diet decreases the allogenicity of donor CD4^+^ T cells in the large intestine.

**Figure 6.**
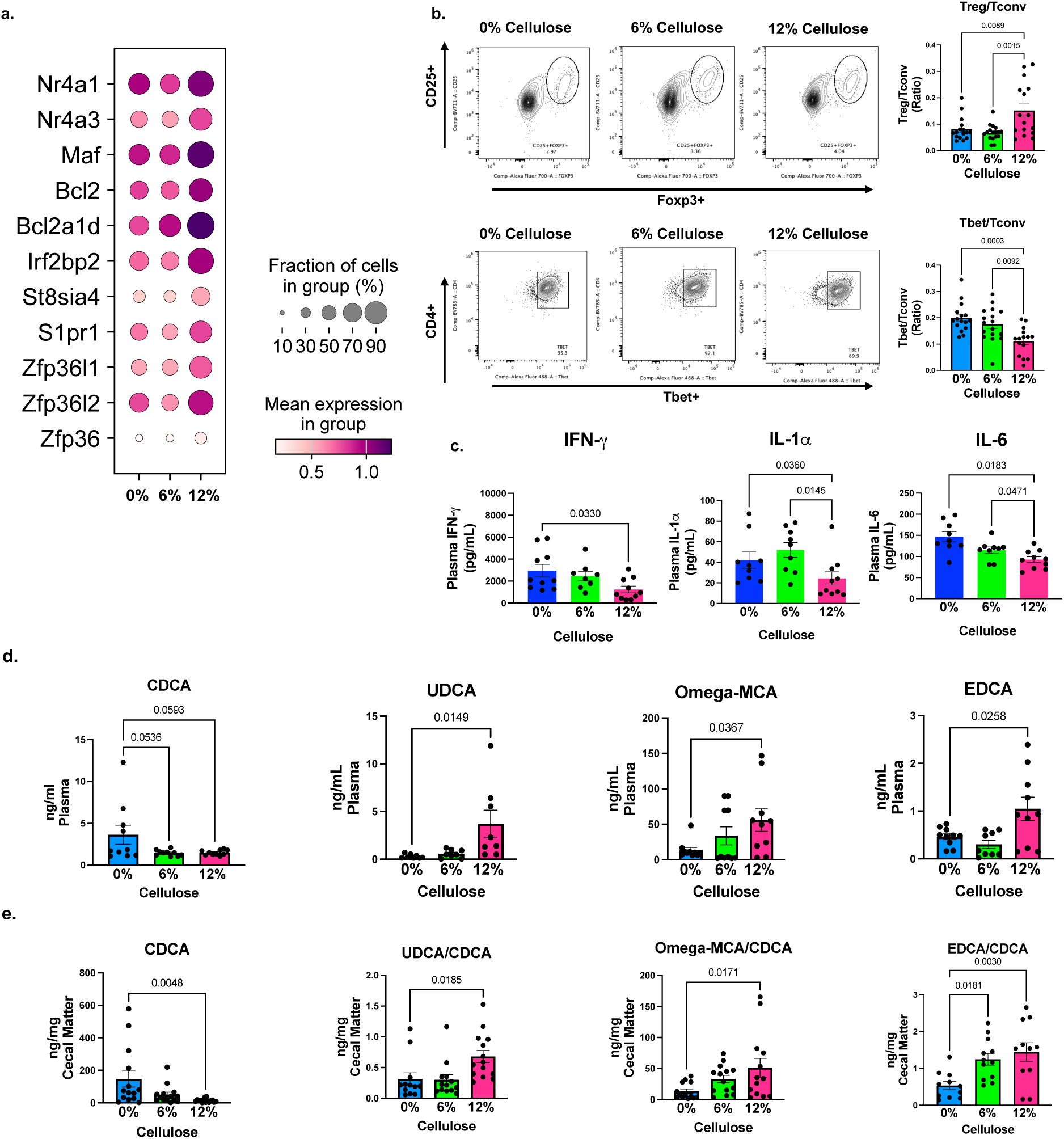
A 12% cellulose diet reduces systemic inflammation in GVHD mice and enriches bile acid transport pathways in CD4⁺ T cells. a: Dot plot of gene expression in CD4⁺ T cells from the large intestine, showing the expression of activation/proliferation and inflammatory genes in cellulose-treated mice. Single cell, 10X. GVHD mice, day 7 post BMT, treated with 0%, 6% and 12% cellulose (n=3 per group). b: Frequency of Tbet⁺ CD4⁺ T cells (Tbet/Tconv ratio). GVHD mice, day 7 post BMT, treated with 0%, 6% and 12% cellulose. Flow cytometry. N=45 (n=15 per group), from 3 independent experiments. One-Way ANOVA followed by Tukey’s test. c: Plasma cytokine concentrations of IFN-γ, IL-1α, and IL-6 levels compared to across fiber diets. N=45 (n=15 per group), from 3 independent experiments. One-Way ANOVA followed by Tukey’s test. d-e: Plasma bile acid concentrations (d) and cecal bile acid concentrations (e) from mice treated with 0%, 6%, and 12% cellulose. Plasma and cecum were collected at day 7 post-allo-HCT. LCMS analysis, n=15 per group. One-Way ANOVA followed by Tukey’s test.

**Supplemental Figure 6:**
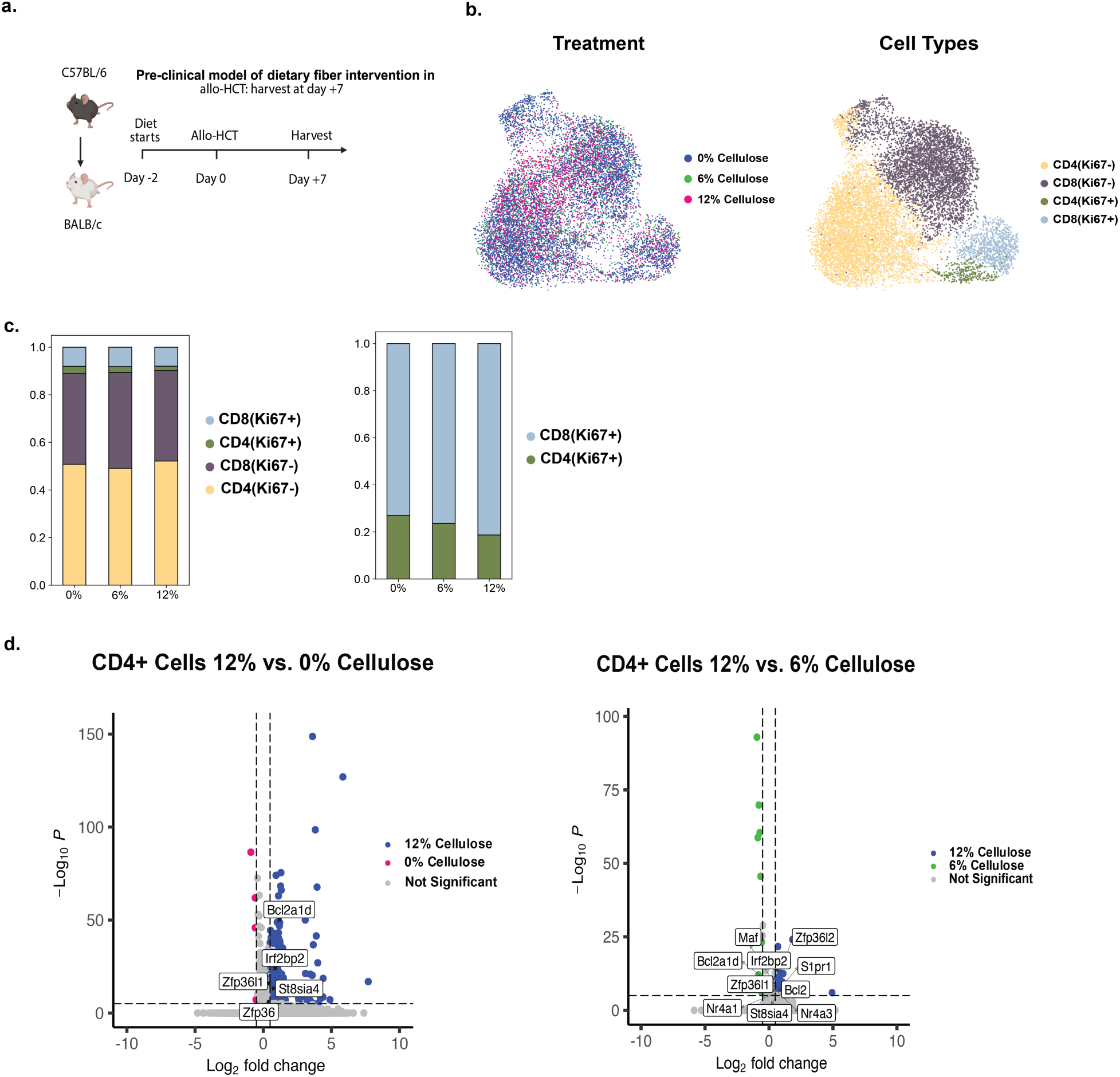
Single-cell transcriptomic analysis reveals differential T cell frequencies and gene expression in the large intestine of allo-HCT mice fed a 12% cellulose diet compared to lower-cellulose diets. **a:** Experimental design for harvesting of large intestine for single-cell analysis, 10X (n=3 per group). **b:** UMAP projections depict immune cell populations from the large intestine of mice receiving diets containing 0%, 6%, or 12% cellulose. Cells are annotated by diet group and by T-cell subset and proliferative status (CD4⁺Ki67⁺, CD4⁺Ki67⁻, CD8⁺Ki67⁺, CD8⁺Ki67⁻). **c:** Bar plots summarize the relative proportions of CD4⁺ and CD8⁺ T cells and the fraction of Ki67⁺ proliferating cells within each dietary group. **d:** Volcano plots display differential gene expression in CD4⁺ T cells comparing 12% cellulose to 0% and 6% cellulose diets, with thresholds for statistical significance and fold-change indicated.

To confirm these findings through protein analysis, we harvested large intestines and plasma from GVHD mice at day +7 post-transplant for flow cytometry and cytokine analysis. Based on prior studies showing that microbial-derived butyrate can promote Treg cell differentiation and suppress GVHD severity^19,37,73^, we quantified the ratio of Treg cells (CD4⁺CD25⁺FOXP3⁺) to conventional CD4⁺ T cells (CD4^+^CD25^+^FOXP3^−^) (Treg/Tconv). Mice fed a 12% cellulose diet had significantly higher Treg/Tconv ratios compared to those fed 0% (p=0.009) and 6% (p=0.002) cellulose diets (**Fig. 6b**), consistent with an anti-inflammatory T cell response. No significant differences were observed between the 0% and 6% cellulose groups. In contrast, the 12% cellulose-fed mice had a significantly lower proportion of proinflammatory Tbet⁺ CD4⁺ cells, compared to the 0% (p<0.001) and 6% (p=0.009) cellulose groups, confirming a shift away from Th1-type inflammation.

To assess systemic inflammation, we used a multiplex flow cytometry assay to measure plasma levels of key cytokines at 7 days post-transplant. Mice fed 12% cellulose showed significantly lower concentrations of IFN-γ (p=0.033 vs. 0%), IL-1α (p=0.036 vs. 0%, p=0.015 vs. 6%), and IL-6 (p=0.018 vs. 0%, p=0.047 vs. 6%) compared to the lower-fiber groups (**Fig. 6c**). These findings are consistent with dampened immune activation, providing evidence that a high-cellulose diet is anti-inflammatory. Together, these data demonstrate that a 12% cellulose diet reduced effector T cell activation, promoted Treg differentiation, and decreased systemic inflammation in GVHD mice.

### A 12% cellulose diet increases immunoregulatory secondary bile acids

Bile acids (BAs) facilitate the absorption of lipids in the small intestine but also influence many biologic functions, including modulating immune signaling^74,75,76^. In addition, secondary BAs, derived from cholesterol metabolism by the microbiota (i.e., *Ruminococcaceae*^74^) have anti-inflammatory properties in the intestine^22,74,75,76,77^. We investigated the effect of dietary cellulose on the BA pool by quantifying plasma and cecal BAs.

In mice fed a 12% cellulose diet, we observed a trend toward reduced plasma concentrations of the primary BA chenodeoxycholic acid (CDCA) (**Fig. 6d**), a known agonist of the farnesoid X receptor (FXR) receptor that is associated with proinflammatory processes and exacerbation of T-cell activation^22,78^. These mice also had higher plasma concentrations of several secondary BAs, including ursodeoxycholic acid (UDCA) (p=0.015), omega-muricholic acid (MCA) (p=0.037), and 3-epideoxycholic acid (EDCA) (p=0.026), compared with mice fed diets of lower cellulose concentrations.

We further analyzed cecal BA composition to determine whether the 12% cellulose diet enhanced microbial conversion of primary to secondary BAs. Mice fed 12% cellulose had significantly lower cecal concentrations of the primary BA CDCA (p=0.0048), and correspondingly significantly elevated ratios of secondary/primary BAs, including UDCA/CDCA, omega-MCA/CDCA, and EDCA/CDCA, (p=0.019, p=0.017, and p=0.0030, respectively), compared with mice fed a 0% cellulose diet (**Fig. 6e**). These findings suggest that high cellulose intake promotes a microbiome capable of more extensive BA metabolism, while promoting a BA pool enriched in immunoregulatory secondary BAs.

## Discussion

In this study, we integrated prospectively collected dietary intake data with longitudinal microbiome and metabolite profiling, providing evidence that dietary fiber intake is associated with clinical outcomes in allo-HCT. The peri-transplant hospitalization period offers a controlled clinical environment that enables precise dietary assessment and longitudinal microbiome sampling, providing a unique opportunity to study these relationships. To our knowledge, this study provides the most comprehensive assessment of patient dietary intake with metabolic readouts in a cohort in this clinical setting.

We found that higher pre-transplant fiber intake correlated with improved OS and reduced incidence of severe acute GVHD. These associations were accompanied by greater microbial diversity; enrichment of butyrate-producing taxa such as *Blautia*, associated with reduced GVHD-related mortality following allo-HCT^15^; and reduced abundance of *Enterococcus,* an opportunistic pathogen linked to poorer OS in allo-HCT recipients^16^. Patients who developed LGI-GVHD exhibited marked depletion of fecal SCFAs, including butyrate and acetate, which are linked to immunoregulatory and protection of the intestinal epithelium. In a complementary preclinical allo-HCT model, we found that cellulose supplementation was associated with improved survival, preserved epithelial integrity, increased microbial diversity, and elevated cecal butyrate concentrations. The cellulose-rich diet also enhanced microbial pathways involved in complex carbohydrate fermentation, SCFA production, and micronutrient synthesis, supporting a mechanistic link between fiber intake, microbiome composition, and immune regulation.

Our findings extend prior observations that microbiome diversity, butyrate-producing taxa, and microbial metabolites, including SCFAs and BAs, are associated with favorable transplant outcomes^15,16,27,33,37,38,77,79^. These results are consistent with reports from the American Gut Project linking fiber consumption and plant diversity to higher α-diversity and SCFA-producing bacteria^80^, as well as feasibility trials of resistant starch intervention for allo-HCT patients^40^, which further support the role of dietary fiber in shaping microbiome composition and metabolite profiles.

In our preclinical allo-HCT model, we found that a cellulose-rich diet is associated with reduced epithelial permeability and increased expression of genes involved in immune tolerance and regulation in allogenic CD4⁺ T cells, aligning with previous reports that microbial metabolites such as SCFAs and BAs reinforce epithelial barrier integrity and regulate T-cell^17,18,19,20,21,81^. Plasma and cecal metabolite profiling revealed a shift from the primary bile acid CDCA, a known FXR agonist associated with the worsening of GVHD, to bile acids like UDCA, which inhibits T-cell activation and improves allo^22,77^. We have previously demonstrated that allo-HCT patients treated with UDCA have lower risk of GVHD-related mortality, indicating the therapeutic potential of using UDCA to restore microbial BA metabolism and mitigate GVHD^77^. The markedly reduced fecal concentrations of butyrate and other SCFAs in LGI-GVHD patients reinforces our hypothesis on the link between microbial metabolite depletion and disease severity. Given that cellulose-degrading bacteria are not well supported by modern diets such that microbiota fiber fermentation and production may be limited^42^, clinical fiber supplementation may be important. Our study and others underscore the importance of dietary components in modulating microbiome-mediated immune responses.

Collectively, our human and murine data show that dietary fiber enhances microbial diversity, supports SCFA production, and maintains epithelial integrity, establishing a strong rationale for fiber-supplementation interventional trials to reduce GVHD and improve allo-HCT outcomes. Our rigorous daily diet monitoring revealed reductions in fiber intake post-transplantation, which may reflect complications and chronic disease that disrupt gastrointestinal function^82,83^. In addition, the median fiber intake in our cohort (7.7 g/day pre-transplant, 4.5 g/day post-transplant) was substantially below recommended levels (25–30 g/day) and U.S. averages (16 g/day)^84,85^, highlighting that a fiber intervention may be particularly needed in this clinical setting and potentially for other hospitalized patients undergoing intensive therapeutic interventions. Overall, our findings demonstrate proof-of-concept that dietary intake during an active cancer therapy is a clinically meaningful determinant of outcomes, with relevance to transplantation, oncology, supportive care, and microbiome biology. Future studies should confirm the importance of dietary fiber through a randomized controlled trial and define optimal fiber types, timing, and integration with supportive care to enhance microbiome-mediated immune regulation.

## Methods

### Patients

Recipients of allo-HCT who were treated by the Adult Bone Marrow Transplantation Service at Memorial Sloan Kettering Cancer Center (MSKCC) and had been consented to an IRB-approved biospecimen collection and diet monitoring were followed in this study.

### Dietary collection and annotation

Dietary data were collected as described in Dai, 2024^29^. Briefly, patients received each meal from the hospital cafeteria, prepared according to standardized recipes and portions, accompanied by a printout to record their food consumption for that meal. Participants were asked to record immediately after each meal whether they consumed 0%, 25%, 50%, 75% or 100% of each item ordered for that meal. A research dietitian or research assistant visited each patient 3 times weekly to verify the meal portion entries and perform 24 h recall to complete any missing entries. If any entries were missing or unable to be verified, the research team annotated the data to ensure exclusion from the final nutrient dataset. Patients were also provided a log to report consumption of any other foods not served through the hospital cafeteria system. Food intake values were verified by the research dietitian and consolidated with the Food and Nutrient Database for Dietary Studies (FNDDS). To prepare the dataset for analysis, water fractions from the FNDDS dataset were used to compute the dehydrated weight of consumed food by converting the volume of food to grams based on 1.05 g/mL. The grams of FNDDS food items consumed in each meal and on each day of hospitalization were listed, enabling accurate comparison of nutrient breakdowns across different food types. The ingredients and their proportions in meals were annotated using *Computrition* software and the USDA FoodData Central website^84^ to identify the main fiber classification. Food types classified as fiber-rich carbohydrates were further annotated based on fiber type (soluble or insoluble) and subtype composition (resistant starch, pectin, lignin, inulin, hemicellulose, beta-glucan, and cellulose) by a research dietitian.

### Statistical analysis for clinical data

Multivariable Cox proportional hazards regression models were fitted for OS for all patients with available pre-transplant dietary data. The hazard of death was assumed to be a function of the average natural logarithm (ln) transformed grams of macronutrient intake and days of broad-spectrum antibiotic exposure during a pre-transplant window (day -7 to day -1), disease type (AML versus others), conditioning intensity, and graft source (T-cell depleted [TCD] versus non-TCD; in which donor T cells are retained in the graft and GVHD risk is significantly higher than with TCD grafts)^33^. Multivariable Fine-Gray proportional subdistributional hazards regression models were fitted for severe (grade 2 or above) acute GVHD and severe acute lower-GI (LGI) GVHD for non-TCD patients with available pre-transplant dietary data. Relapse and death were regarded as competing risks of severe acute GVHD, while relapse, death, and severe acute GVHD at other organs were regarded as competing risks of severe acute LGI GVHD. The same covariates as used in the OS models, except graft source, were also used for the GVHD models. Cox and Fine-Gray models were fitted for the five macronutrients (fiber, sugar, fat, protein, and other carbohydrates). P-values for the hazard ratios were adjusted across macronutrients by the Benjamini-Hochberg method. OS probability and severe acute GVHD and LGI-GVHD cumulative incidence functions were respectively visualized by Kaplan-Meier and Aalen-Johansen curves^86,87^, with patient groups defined by higher or lower than the median average grams of pre-transplant dietary fiber intake. Landmark analysis for OS and severe acute GVHD, landmarked at day 30 after transplant using similar Cox and Fine-Gray models was also performed as described above. This analysis included the average post-transplant (day 0 to day +30) ln-transformed grams of intake in the model and extended the window used to calculate the duration of broad-spectrum antibiotic exposure from day -7 to day 30. Associations were reported between post-transplant average nutrient intakes and day 30-landmarked clinical outcomes. Significance was defined as an FDR-adjusted p-value < 0.05.

### Fecal microbiome analysis by 16S rRNA and shotgun sequencing

Bacterial cell walls were disrupted and nucleic acids isolated using silica bead-beating and phenol-chloroform extraction, and the V4-V5 variable region of the 16S rRNA gene was amplified. Amplicons were purified either using a Qiagen PCR Purification Kit (Qiagen) or AMPure magnetic beads (Beckman Coulter) and quantified using a Tapestation instrument (Agilent). DNA was pooled to equal final concentrations for each sample and then sequenced on the Illumina platform. 16S sequencing data were analyzed using the R package DADA2 (version 1.16.0) pipeline with default parameters except for maxEE=2 and truncQ=2 in filterandtrim function. Amplicon sequence variants (ASVs) were annotated according to the NCBI 16S database using BLAST. Shotgun sequencing reads were taxonomically profiled using MetaPhlAn3 (MetaPhlAn 4.1 for mouse metagenomics data). These reads and profiles were then passed through HUMAnN3 (HUMAnN 3.9 for mouse), a pipeline for profiling the presence and abundance of microbial pathways. CAZyme abundances were computed using run_dbcan v4.2.0-rc.1. Microbiome α-diversity was evaluated using the inverse Simpson index, a summary statistic of both the richness and evenness of the bacterial community. Taxa abundances were summarized at the genus level.

### Statistical analysis of microbiome data

For clinical 16S sequencing data, generalized estimating equations (GEE) were used to model the mean trajectory of day 0 to day 30 log-transformed longitudinal inverse Simpson index as a function of cubic spline basis functions of time, average log-transformed grams of dietary intake, and days of broad-spectrum antibiotics exposure during pre-transplant (day -7 to 0) or peri-transplant (day -7 to 30) windows. An exchangeable working correlation structure was assumed across repeated measurements. Similar GEE models were also used to model centered log-ratio transformed *Blautia* and *Enterococcus* abundance. The above GEE models were applied for the 7 common fiber subtypes (resistant starch, pectin, lignin, inulin, hemicellulose, beta-glucan, and cellulose). Wald p-values with respect to the dietary intake effects were adjusted for multiple testing by the Benjamini-Hochberg approach. Significance was defined as a false discovery rate (FDR)-adjusted p-value < 0.05.

Mouse shotgun sequencing data were analyzed by redundancy analysis (RDA) using Euclidean distance on Hellinger transformed species abundances with fiber as a discrete covariate and experiment batch partialled out. Covariate significance was tested in the RDA model using permutational ANOVA with 9999 permutations. FLORAL was used to compute pathway selection probabilities with fiber % as a continuous outcome using multivariable compositional lasso models ^47^; five-fold cross-validation was repeated 100 times to obtain the average effect sizes and selection probabilities of the selected pathways. P-values for testing associations between cellulose % *vs.* taxon-specific relative abundances, α-diversity, or CAZyme abundance were computed using Gamma generalized linear models with sample batch included as a covariate. P-values were adjusted using the Benjamini-Hochberg method. Significance was defined as an FDR-adjusted p-value < 0.05.

### Gas chromatography-mass spectrometry

Gas chromatography-mass spectrometry (GC-MS) for fecal SCFA concentrations was performed on samples from a sub-cohort of patients with acute lower-GI GVHD or without GVHD, matched by stool sample availability. ***Fecal sample preparation*:** Fecal samples (∼120 mg) were weighed into 2 mL microtubes containing 1.4 mm ceramic beads (Omni International) and resuspended to a final concentration of 100 mg/mL using 80:20 methanol-to-water containing acetate-d3, propionate-d5, butyrate-d7, and valerate-d9 as internal standards (Cambridge Isotope Laboratories). Fecal samples were homogenized using a Bead Ruptor (Omni International) at 6 m/s for 3 min at 4°C, then centrifuged for 20 min at 20,000 x *g* at 4°C. Sample extracts (100 µL) were added to 100 µL of 100 mM borate buffer (pH 10), and extracted with 400 µL of 100 mM pentafluorobenzyl bromide derivatization reagent (Thermo Scientific) diluted in acetonitrile and 400 µL of cyclohexane. Samples were analyzed at 1:100 dilution (prepared with cyclohexane). Calibration curve levels were prepared in borate buffer, ranging from 0.1 to 100 mM. For quality control samples (QCs), three SCFA concentrations (0.04, 0.4, and 4 mM) were spiked into murine SCFA-depleted cecal control samples. ***GC-MS analysis and SCFA quantification:*** GC-MS analysis used an Agilent 7890A GC and Agilent 5975C MS detector operating in negative chemical ionization (nCI) mode. Methane was used as the chemical ionization reagent gas at 2 mL/min, and a 1 µL splitless injection was made onto a VF-1701ms column (30 m x 0.25 mm, 0.25 µm; Agilent Technologies). Data analysis was performed using Agilent MassHunter Quantitative Analysis software (Version 10.1, Agilent Technologies).

### Statistical analysis of GC-MS data

GEEs were applied to model the mean trajectory of day 0 to day 30 longitudinal SCFA concentrations as a function of cubic spline basis functions of time, batch of GC-MS assays, average log-transformed grams of dietary intake and days of broad-spectrum antibiotic exposure during pre-transplant (day -7 to 0) or peri-transplant (day -7 to 30) windows. An exchangeable working correlation structure was assumed across repeated measurements. The above GEE models were applied for the 7 SCFAs (2-methylbutyrate, acetate, butyrate, isobutyrate, isovaleric acid, propionate, and valeric acid) for three selected fiber subtypes (resistant starch, hemicellulose, and cellulose). P-values associated with fiber intake were adjusted across fiber subtypes by the Benjamini-Hochberg method for models corresponding to each type of SCFA. Significance was defined as an FDR-adjusted p-value < 0.05.

### Differential expression analysis

was performed using Scanpy’s rank_gene_groups^88^, function with default parameters and using the 2-sided Wilcoxon rank-sum method (X). Significance was defined as an FDR-adjusted p-value < 0.05.

### Pathway enrichment analysis

was performed with GSEA (v4.3.2) 30 as per gene list and rank metric provided. The GSEA Preranked module was used to predict pathway enrichment in threshold free comparisons: (a) 12% cellulose vs. 6% cellulose, (b) 12% cellulose vs. 0% cellulose, (c) 0% cellulose vs. 6% cellulose. Rankings were created for differentially expressed genes using the Wilcoxon Z-score in descending order. Predicted pathways with an FDR ≤ 0.05 were considered significantly enriched^88^.

### Statistical analysis for murine experiments

Analysis of processed data was conducted using R 4.4.2, Graphpad Prism v7.5.0. Flow cytometry data were analyzed using FlowJo™ v10.8 Software (BD Life Sciences). Group sizes for murine BM+T experiments were based upon the statistical premise that 30 animals are required to detect a 50% difference with statistical significance. Statistical significance was established as p-value < 0.05. For comparisons between mice receiving BM vs BM+T cells, a two-sided Wilcoxon rank-sum test was used. Gene expression analysis was conducted using a Wilcoxon rank-sum test. Significance was defined as an FDR-adjusted p-value < 0.05.

### Cell isolation from murine tissues for single-cell sequencing and flow cytometry

Colon samples were processed by removing adherent adipose tissue and resecting Peyer’s patches. Colons were opened longitudinally and shaken vigorously in 1X PBS to release contents. Tissues were incubated in 25 mL intestinal intraepithelial lymphocyte (IEL) solution (1× PBS with 2% FBS [ThermoFisher], 10 mM HEPES buffer [ThermoFisher], 1% penicillin/streptomycin [ThermoFisher], 1% L-glutamine [ThermoFisher], 1 mM EDTA [Sigma], and 1 mM dithiothreitol [DTT; Sigma] added immediately before use) for 15 min at 37 °C with vigorous shaking (250 rpm). Tissues were then removed from IEL suspension, rinsed in PBS, and transferred into 50 mL tubes containing 3 one-quarter-inch ceramic beads (MP Biomedicals) and 25 mL collagenase solution (1× RPMI 1640 with 2% FBS [ThermoFisher], 10 mM HEPES buffer [ThermoFisher], 1% penicillin/streptomycin [ThermoFisher], 1% L-glutamine [ThermoFisher], 1 mg/mL collagenase D [Sigma], and 1 U/mL DNase I [Sigma]). Following incubation for 30 min at 37 °C with vigorous shaking (250 rpm), digested lamina propria samples were passed through a 100-μm strainer and washed to remove collagenase with washing solution (1× RPMI 1640 [w/o HEPES; ThermoFisher], 5% FCS [ThermoFisher], 10 mM HEPES [Sigma], and 1% L-glutamine [ThermoFisher]). Tissues were washed and centrifuged twice (5 min, 1200 RPM), resuspended, and counted.

### Flow cytometry of colon tissue from murine models

Cells were stained with Zombie UV (Biolegend) in PBS at room temperature for 30 min. Cells were subsequently stained with Fc-block for 20 min, followed by fluorescent surface antibodies for 20 to 30 min, then fixed and permeabilized with FoxP3 Fixation/Permeabilization kit (Invitrogen), and stained with intracellular antibodies. Staining was performed in PBS + 0.5% BSA + 2% FBS or Brilliant Violet Stain Buffer (BD) unless otherwise indicated. After staining, cells were washed with and resuspended for analysis in PBS + 0.5% BSA + 2% FBS. Data were collected using a Cytek Aurora cytometer.

The following anti-mouse antibodies were used: CD45-BUV395 (clone 30-F11, BD Biosciences), CD3-APC-Cy7 (clone 145-2C11, BD Biosciences), CD4-BV785 (clone RM4-5, BioLegend), CD8-PE-Cy7 (Clone 53-6.7, BD Biosciences), H-2kD-PE (clone SF1-1.1, BioLegend), H-2kB-BUV805 (clone AF6-88.5, BD Biosciences), Gr1-PE-Cy5 (clone RB6-8C5, ThermoFisher), CD11b PE-Cy5 (clone M1/70, ThermoFisher), Ter119-PE-Cy5 (clone TER-119, BioLegend), CD11c PE-Cy5 (clone N418, ThermoFisher), RORgt-PE-CF-594 (clone Q31-378, BD Horizon), Tbet-Alexa 488/FITC (clone 4B10, Santa Cruz Biotech), Foxp3-Alexa Fluor 700 (clone FJK-16s, eBioscience), Ki67-Pacific Blue (clone 16A8, BioLegend), EpCAM PE-Cy7 (clone G8.8, eBioscience) and MHC-II-APC (clone M5/114.15.2, eBioscience).

### Cytokine quantification in plasma

Mouse plasma was collected at day 7 post allo-HCT and stored at −80 °C until processing. Each sample was analyzed using the LEGENDPlex (Biolegend) beads-based assay, following the manufacturer’s protocol. The Mouse Inflammation Panel was used with the following targets: IL-1α, IL-1β, IL-6, IL-10, IL-12p70, IL-17A, IL-23, IL-27, CCL2 (MCP-1), IFN-β, IFN-γ, TNF-α, and GM-CSF. Data were collected using a Penteon Agilent cytometer.

### Single-cell RNA sequencing

Murine large intestine cells were collected for single-cell analysis as described above. In brief, murine large intestine cells were washed once with PBS containing 1% BSA and resuspended in PBS containing 1% BSA to a final concentration of 1000 cells/µL. Individual cell suspensions were incubated for 30 min with hashtag oligonucleotide-conjugated antibodies. ***Hashtag antibodies*:** TotalSeqTM-B0301 anti-mouse Hashtag 1 (Cat#155831; 1:50); TotalSeqTM-B0301 anti-mouse Hashtag 2 (Cat#155833; 1:50); TotalSeqTM-B0301 anti-mouse Hashtag 3 (Cat#155835; 1:50); TotalSeqTM-B0301 anti-mouse Hashtag 4 (Cat#155837; 1:50).

### Single-cell transcriptome sequencing

Suspension viability and cell number of single-cell suspensions were assessed using Trypan blue staining and a Countess II Automated Cell Counter (ThermoFisher). Cells were then loaded onto a Chromium Next GEM Chip G (10x Genomics PN 1000120) for GEM generation, cDNA synthesis, cDNA amplification, and library preparation of ∼17–63,000 cells using the Chromium Next GEM Single Cell 3’ Kit v3.1 (10x Genomics PN 1000268) according to the manufacturer’s protocol. cDNA amplification included 11–12 cycles; 10 µL of the material was used to prepare sequencing libraries via 14 cycles of PCR. Indexed libraries were pooled by molarity and sequenced on a NovaSeq X in a PE28/88 paired end run using the NovaSeq X 10B Reagent Kit (Illumina). An average of 23,000 paired reads were generated per cell.

### Cell surface protein feature barcode analysis

Amplification products generated using the methods described above included both cDNA and feature barcodes tagged with cell barcodes and unique molecular identifiers. Smaller feature barcode fragments were separated from longer amplified cDNA through a 0.6X cleanup using aMPure XP beads (Beckman Coulter catalog # A63882). Libraries were constructed using the 3’ Feature Barcode Kit (10X Genomics PN 1000276) according to the manufacturer’s protocol with 10 cycles of PCR. Indexed libraries were pooled equimolar and sequenced on a NovaSeq X in a PE28/88 run using the NovaSeq X 10B Reagent Kit (Illumina). An average of 325 million paired reads were generated per sample.

### Single-cell preprocessing and quality control

Raw FASTQ files were processed with Cell Ranger (v7.0.0) to generate gene-barcode matrices and mapped to the mm10-2020-A mouse reference. 0%, 6%, and 12% cellulose conditions were processed and analyzed as separate datasets through normalization, dimensionality reduction, and clustering. Cluster-level results were subsequently compared across conditions after removal of contaminating CD45-cell populations. The filtered_feature_bc_matrix.h5 files were imported into the shunPykeR workflow^88^. Subsequently, genes not detected in any cell as well as ribosomal and hemoglobin genes were removed, and counts were normalized to 10,000 reads per cell and log-transformed with a pseudo-count of 1. Dimensionality reduction was performed by principal component analysis (PCA), followed by Leiden clustering. Cell quality was evaluated using total unique molecular identifier (UMI) counts, number of detected genes, and mitochondrial and ribosomal read fractions. Cells with low complexity (≤1000 genes), low counts, and/or high mitochondrial content (≥20%) were flagged as poor quality, and clusters enriched for such cells were excluded. Doublets were identified with scrublet and removed. Cells expressing epithelial (Epcam+) or myeloid (Itgam)+ markers were also filtered out as non-target contaminants.

### Single-cell data integration

After quality control and initial annotation, GHVD datasets for the 0% plus 6% and 12% cellulose groups were concatenated. PCA and clustering were then repeated, and major cell types were annotated by canonical marker expression. Residual epithelial contamination was minimized by computing epithelial scores using scanpy’s sc.tl.score_genes^89^ with a predefined epithelial gene set. Such cells with epithelial scores >0 were removed. PCA and clustering were rerun to obtain the final integrated dataset for downstream analyses.

### Differential gene expression analysis

Differential expression between samples in the cell types of interest were computed with scanpy’s sc.tl.rank_gene_groups, using the Wilcoxon rank-sum test. Genes were considered differentially expressed at an FDR-adjusted p-value ≤ 0.05, without applying an additional fold-change cutoff.

### Bile acid analyses

Bile acid analyses of murine plasma and cecal samples were performed using stable isotope dilution as previously described^77^. Briefly, plasma bile acids were extracted using 10-fold excess acetonitrile : water : methanol : formic acid (7:2:1:0.02, v/v/v/v) spiked with 20 ng/mL of internal standards (d4-CA, d4-CDCA, d4-ω-MCA, d4-CDCA, d4-UDCA, d4-EDCA, d4-LCA, d4-IALCA; Cayman Chemical). Samples were vortexed and incubated at −80 °C for protein precipitation, followed by centrifugation (20,000 x *g*, 20 min, 4°C). The supernatant was dried under vacuum centrifugation and reconstituted in methanol: 10 mM ammonium acetate in water (1:1, v/v) for LCMS analysis. For cecal samples, 80 mg of cecal content was resuspended in 500 µL of extraction solvent (methanol: water, 4:1, v/v) spiked with internal standards (50 ng/mL). The samples were homogenized using bead beater (Benchmark Scientific) for 3 cycles, each with 30 s bead beating at 420 rpm, and 30 s of pause. The homogenate was clarified and concentrated as described above. The dried concentrate was reconstituted in 2-fold excess volume of water: methanol (1:1, v/v) for bile acid analysis. Calibration curves for bile acids (CA, CDCA, MCA, CDCA, UDCA, EDCA, LCA, IALCA; Cayman Chemical) were prepared in 3.5% BSA (w/v) over linear ranges of 0–616 µM primary bile acids and 0–32 µM for secondary bile acids, respectively. The LCMS analysis was performed using Vanquish Flex UHPLC coupled to TSQ-Altis Plus mass spectrometer (ThermoScienfic). The bile acids were separated on a BEH C18 (2.1x 50 mm; Waters) column at a 0.5 mL/min flow rate using solvent A (5 mM ammonium acetate and 0.1% formic acid in water) and solvent B (5 mM ammonium acetate and 0.1% formic acid in 95% acetonitrile), and a 25 min run time (0.5 min, 5% B; 8.9 min, 5-40% B; 4 min, 40–55% B; 4 min, 55–59.3% B; 1 min, 59.3–95% B; 3 min, 95% B, 1 min, 95–5% B; 3 min at 5% B). The column temperature was kept at 65 °C. The bile acids were analyzed using intact precursor ions as product ion masses in negative mode. All bile acids exhibited excellent linearity (R2 >0.9) with limit of detections ranging from 2–92 fmol on column and precision <15% relative standard deviation (RSD) and accuracy >85% for bile acid concentrations above the limit of quantitation.

### Mice

Female C57BL/6 mice, aged 6–8 weeks, were obtained from Jackson Laboratory. Fecal samples collected during transplantation course included pre-dietary intervention, day –2 (start of dietary intervention), and days 0, +7 and +14 relative to allo-HCT. All mouse experiments were conducted with approval of the Institutional Animal Care and Use Committee at MSKCC (protocols #23-03-006 and #99-07-025).

### Mouse diet and experimental groups

Mice were assigned to experimental diets of differing fiber composition (0% [fiber-free], 6% cellulose [comparable to standard chow], and 12% cellulose [high fiber]) while maintaining similar concentrations of other macro-nutrients. Mice were monitored for survival for 90 days post-transplant. Cohorts included BM only (n=15 per group) and BM+T (n=30 per group), for a total of 135 mice from 3 independent experiments.

### GVHD mouse model

C57BL/6 (donors) and BALB/c mice (recipients) were procured from the Jackson Laboratory, housed 5 mice/cage under standard specific-pathogen-free conditions (chow and water ad libitum, 12 h light cycle). BMT was performed as previously described^90^. Briefly, following split-dose lethal irradiation with 900 cGy, mice received 5 × 10^6^ bone marrow cells that were depleted of T cells anti-Thy-1.2 (BioXcell) and Low-Tox-M rabbit complement (CEDERLANE Laboratories), and 10^6^ magnetic-bead-purified splenic T cells via anesthetized retro-orbital injection. Mice were monitored daily for survival and weekly for GVHD clinical scores, as described previously^90^.

## Acknowledgements

We acknowledge the use of the Integrated Genomics Operation Core, at Memorial Sloan Kettering Cancer Center, funded by the NCI Cancer Center Support Grant (CCSG, P30 CA08748), Cycle for Survival, and the Marie-Josée and Henry R. Kravis Center for Molecular Oncology, Hannah Rice and Keely Walker for editorial support.

---

**Supplemental Table 1.** Summary of daily dietary intake of fiber, fiber subtypes, and macro-nutrients pre-transplant (days -7 to -1, grams per day)

| Characteristic | N = 171<sup>1</sup> |
| --- | --- |
| **Fiber** | 7.7 (5.0, 12.2) |
| Soluble fiber | 4.77 (2.97, 7.10) |
| Insoluble fiber | 2.10 (1.34, 3.93) |
| Cellulose | 0.71 (0.24, 1.61) |
| Beta glucans | 0.18 (0.00, 0.74) |
| Hemicellulose | 0.90 (0.25, 1.58) |
| Inulin | 0.15 (0.00, 0.72) |
| Lignin | 0.23 (0.00, 0.62) |
| Pectin | 0.82 (0.45, 1.72) |
| Resistant starch | 1.82 (1.15, 3.15) |
| **Sugar** | 70 (47, 96) |
| **Fat** | 39 (21, 60) |
| **Protein** | 38 (20, 50) |
| **Carbohydrates** (excluding fiber and sugar) | 67 (38, 99) |
| **Total weight** | 237 (153, 321) |

<sup>1</sup>Median (Q1, Q3)

**Supplemental Table 2.** Summary of daily dietary intake of fiber, fiber subtypes, and macro-nutrients peri-transplant (days -7 to 30, grams per day)

| Characteristic | N = 171<sup>†</sup> |
| --- | --- |
| **Fiber** | 4.5 (2.6, 7.8) |
| Soluble fiber | 3.00 (1.78, 4.87) |
| Insoluble fiber | 1.09 (0.45, 2.24) |
| Cellulose | 0.33 (0.10, 0.92) |
| Beta glucans | 0.18 (0.05, 0.52) |
| Hemicellulose | 0.31 (0.13, 0.74) |
| Inulin | 0.18 (0.02, 0.46) |
| Lignin | 0.15 (0.06, 0.32) |
| Pectin | 0.72 (0.36, 1.25) |
| Resistant starch | 0.97 (0.52, 1.90) |
| **Sugar** | 57 (38, 75) |
| **Fat** | 21 (12, 42) |
| **Protein** | 22 (12, 41) |
| **Other carbohydrates** (excluding fiber and sugar) | 41 (24, 78) |
| **Total weight** | 156 (101, 257) |

<sup>†</sup>Median (Q1, Q3)

